# Replicative senescence of neural progenitors induces astrocyte senescence in 2D cultures and human midbrain organoids

**DOI:** 10.64898/2026.06.17.732317

**Authors:** Virginia Cora, Daniele Ferrante, Nika Lu-Yang, Javier Jarazo, Alise Zagare, Jens C. Schwamborn, Silvia Bolognin

## Abstract

Astrocyte senescence (astrosenescence) has emerged as a potential mechanism through which ageing may progressively impair glial homeostatic neuronal support and render the brain more vulnerable to neurodegenerative diseases such as Parkinson’s disease (PD). Yet human experimental models that specifically recapitulate astrosenescence within a PD-relevant genetic background are lacking. Here, we applied passage-induced replicative exhaustion to human iPSC-derived neuroepithelial stem cells (NESCs) from a healthy donor and LRRK2-G2019S PD patients to establish an *in vitro* midbrain ageing model. Extended passaging induced senescence-associated changes in NESCs, including reduced proliferative capacity, decreased cleaved caspase 3 (CC3) and telomerase mRNA expression, and DNA damage response (DDR) activation, while preserving differentiation potential. Astrocytes derived from aged progenitors showed an increase in SA-β-gal activity, loss of Lamin B1, DDR activation, and an altered mitochondrial morphology. In human midbrain organoids (hMOs), passage-induced ageing was associated with astrocyte-specific DDR activation and lipidomic remodelling. A multi-modal integrative analysis further revealed that, despite the variable penetrance of individual senescence-associated markers across lines, passage condition constitutes the primary discriminant of the collective cellular state, supporting the view that replicative senescence as a biological program is detectable only when individual readouts are aggregated across modalities. Together, this work establishes a human iPSC-based platform recapitulating key features of cellular senescence in both healthy and LRRK2-G2019S contexts and provides an *in vitro* platform to investigate the contribution of astrosenescence to PD-related neurodegeneration.

## Introduction

Ageing is the strongest risk factor for neurodegenerative diseases such as Parkinson’s disease (PD) ^1,2^. Yet the mechanisms by which age-associated cellular programs render the human midbrain vulnerable to neurodegeneration remain poorly understood. PD is characterised by the progressive loss of dopaminergic neurons in the *substantia nigra pars compacta*, a midbrain region essential for motor control ^3^. While most cases are sporadic, autosomal dominant gain-of-function mutations in LRRK2 (Leucine-Rich Repeat Kinase 2), most notably the G2019S variant, represent the most common cause of autosomal dominant familial PD and are also found in a subset of sporadic cases ^3,4^. PD research has historically focused on neuronal dysfunction, with dopaminergic neurons considered the primary drivers of disease pathogenesis ^5^. However, increasing evidence indicates that non-neuronal cell types, such as astrocytes, critically regulate neuronal survival and shape disease progression ^6^.

Astrocytes are central regulators of neural homeostasis, providing neurotransmitter clearance, reactive oxygen species buffering, metabolic and trophic support, and waste removal ^7^. For these reasons, astrocytes may be especially vulnerable to age-associated changes. Notably, LRRK2-G2019S astrocytes have been shown to exhibit impaired mitochondrial morphology and activity ^8^, altered inflammatory response ^9^, impaired chaperone-mediated autophagy, α-synuclein accumulation and reduced capacity to support neuronal survival ^10^, directly implicating astrocyte dysfunction in LRRK2-associated PD pathogenesis. One mechanism through which astrocytes may progressively lose their homeostatic functions is cellular senescence. During physiological ageing, astrocytes can become senescent, a state termed astrosenescence, characterised by irreversible cell cycle arrest and acquisition of a pro-inflammatory secretory phenotype (SASP) ^11,12^. Accumulation of senescent cells has been implicated in age-associated tissue dysfunction, supported by increased senescent astrocytes observed in PD contexts both *in vitro* and *in vivo* ^13,14^. Astrosenescence has also been shown to negatively affect neuronal survival in mouse and rat models ^14–16^. These findings suggest that astrosenescence may actively contribute to neuronal vulnerability in the ageing brain and can exacerbate PD-associated pathology.

Despite this emerging evidence, the relationship between astrocyte senescence and PD-associated genetic risk within a human midbrain context remains difficult to study. A major limitation is the lack of experimental systems that integrate human genetic background with senescence-associated cellular features. Human-induced pluripotent stem cell (iPSC)-derived models provide access to relevant human cell populations but typically lack features associated with senescence. Existing *in vitro* ageing strategies, including progeria-associated mutations ^17^, telomerase inhibition ^18,19^, exposure to chemical stressors ^20^ or ionising radiation ^21,22^, have shown how senescence can be, at least partially, triggered *in vitro*. However, these methods often rely on genetic or chemical-induced perturbations that may confound disease-relevant phenotypes, especially in the field of PD, which is a complex and multifactorial disease.

Here, we establish a human iPSC-derived midbrain platform that incorporates certain senescence-associated features through passage-induced replicative exhaustion of neuroepithelial stem cells (NESCs). NESC passaging induces molecular changes consistent with cellular senescence, including reduced proliferative capacity, nuclear lamina alterations, and DNA damage response activation (DDR), while preserving differentiation potential. This enables the generation of astrocytes and hMOs enriched for senescence-associated features. Astrocytes derived from high-passage progenitors exhibit DDR activation, reduced proliferative activity, together with apoptosis resistance, and mitochondrial alterations. In aged hMOs, astrocyte-associated senescence features coexist with global lipidomic remodelling. Systematic comparison of category-level senescence effects across NESCs, astrocytes, and hMOs further revealed that the PD genetic background shifts the baseline cellular states toward a senescent-like profile and accelerates the trajectory of senescent-like-associated changes in a category-dependent manner. Together, this work establishes a human midbrain model recapitulating multiple senescence-associated features in a genotype-dependent manner and provides an *in vitro* system to investigate the contribution of astrosenescence to PD-related neurodegeneration.

## Material and Methods

### Data and Code availability

All original and processed data, along with the scripts supporting the findings of this study, are being uploaded to DataVerse, and all the scripts are available here https://github.com/nikaaluu/Aging.git. Gene expression datasets have been deposited in the Gene Expression Omnibus (GEO) under the accession number GSE331395.

### Human iPSCs maintenance

Human iPSC lines used in this study (Supplementary Table 1) were generated and characterised in previous studies ^23–28^. Cells were maintained on Geltrex-coated (Gibco, Cat. No. A1413302) plates at 37°C with 20%O_2_ in E8 medium supplemented with 1× Penicillin/Streptomycin (Gibco, Cat. No.15140122). iPSCs were passaged weekly as single cells following treatment with Accutase (Thermo Fisher, Cat No. A11105-01) for 3-10 minutes at 37°C. Cells were collected, centrifuged at 300xg for 3 minutes, and reseeded at 10,000-50,000 cells/cm^2^ in E8 medium (Gibco, Cat. No. A1517001) supplemented with 5 µM Y-27632ROCK inhibitor (Millipore, Cat. No. SCM075) for 24 hours maximum post-seeding. The generation of the mitophagy reporter sensor iPSC lines (H and PD2) was previously described ^23^.

### Generation and maintenance of human iPSC-derived NESCs

NESCs were generated according to published protocols ^25^. Human iPSCs were harvested as single cells using Accutase (3-10 minutes at 37°C). 1.2 × 10^6^ cells were collected, centrifuged at 300×*g* for 3 minutes, and resuspended in Embryoid Body (EB) medium (KO-DMEM (Gibco, Cat. No. 10829018) supplemented with 10% KnockOut Serum Replacement, 1× Penicillin/Streptomycin, 1× GlutaMax (Gibco, Cat. No. 35050061), 1× NEAA (Gibco, Cat. No. 11140050), 100 µM β-mercaptoethanol (Gibco, Cat. No. 21985023), 3 µM CHIR99021 (Axon Mechem, Cat. No. CT99021), 10 µM dorsomorphin (Tocris, Cat. No. 3093), 0.5 µM PMA (Enzo Life Science, Cat. No. ALX-420-045-M005) supplemented with 10 µM ROCK inhibitor (Y-27632). Cells were seeded in 24-well AggreWell plates (StemCell Technologies, Cat. No. 27845) according to manufacturer instructions to generate Embryoid Bodies (EBs). Plates were centrifuged at 100×*g* for 3 minutes. On day 1, EBs were collected and transferred into an ultra-low attachment 6-well plate in EB medium. On day 2, EB medium was replaced with NESC Patterning medium (DMEM/F-12 (Gibco, Cat. No. 11320033): Neurobasal (Gibco, Cat. No. 21103049) [1:1, v/v] supplemented with 1× B-27 Supplement without Vitamin A (Gibco, Cat. No. 12587010), 1× N-2 Supplement (Gibco, Cat. No. 17502001), 1× Penicillin/Streptomycin, 1× GlutaMax, 3 µM CHIR99021, SB-431542 (Abcam, Cat. No. ab120163), 10 µM dorsomorphin, 0.5 µM PMA). Medium was changed on day 4 of differentiation. On day 6 of differentiation, EBs were mechanically dissociated by pipetting 5-10 times, and the resulting suspension was seeded on Geltrex-coated 6-well plates in NESC Maintenance medium (DMEM/F-12:Neurobasal [1:1, v/v] supplemented with 1× B-27 Supplement without Vitamin A, 1× N-2 Supplement, 1× Penicillin/Streptomycin, 1× GlutaMax, 3 µM CHIR99021, 0.5 µM PMA, Ascorbic Acid (Sigma Aldrich, Cat. No. A4544) to establish NESC cultures.

### Generation and maintenance of human iPSC-derived Neural Stem Cells (NSCs)

After passaging NESCs as described above, the medium was replaced with NSC Maintenance medium I (DMEM/F-12 supplemented with 1× B-27 Supplement, 1× N-2 Supplement, 1× Penicillin/Streptomycin, 20 ng/mL FGF2 (Peprotech, Cat. No. 100-15)). On day 4, cells were passaged using Accutase (5-10 minutes at 37°C), centrifuged at 300×*g* for 3 minutes, and plated onto Geltrex-coated plates in NSC Maintenance medium II (DMEM/F-12 supplemented with 1× B-27 Supplement (Gibco, Cat. No. 17504044), 1× N-2 Supplement, 1× Penicillin/Streptomycin, 20 ng/mL EGF (Peprotech, Cat. No. 100-20), 20 ng/mL FGF2, 10 ng/mL human LIF (Sigma Aldrich, Cat. No. L5283). NSCs were kept in this medium for 3-4 passages, and the medium was changed every 2-3 days.

### Generation of aged and young NESCs and NSCs

To generate aged NSCs or NESCs, cells were categorised based on passage number. NESCs were defined as young at passages <8 and as aged at passages >15. NSCs were defined as young at passages <5 and as aged at passages >13.

### Generation and maintenance of astrocytes

To generate astrocytes, the medium of young and aged NSCs was replaced with astrocyte differentiation medium (DMEM/F-12 supplemented with 1× Penicillin/Streptomycin, 1× L-Glutamine (Tocris, Cat. No. 0218), and 1× FBS (Sigma Aldrich, Cat. No. F7524) ^29^. The medium was changed every 3 days. Between days 18 and 20, cells were passaged using Accutase (3 minutes at 37°C), centrifuged at 300×g for 3 minutes, and seeded onto experimental plates at a density not exceeding 70% confluency. Cells were maintained at 37°C until days 24-25 of differentiation, at which point experiments were performed.

### Generation and maintenance of human iPSC-derived hMOs

hMOs were generated as previously described ^30^. Control or aged NESCs were cultured in Geltrex-coated 6-well plates in NESC Maintenance medium until reaching ∼80% confluence. On day 0, NESCs were dissociated into single cells using Accutase (5-10 minutes at 37°C). Cell suspension was collected, centrifuged at 300×*g* for 3 minutes, and resuspended in fresh NESC Maintenance medium. After cell counting, the cell suspension was diluted to 6 × 10^4^ cells/mL, and 150 µL (9 × 10^3^ total cells) was added to each well of a U-bottom ultra-low attachment 96-well plate (faCellitate, Cat. No. F202003). Plates were maintained at 37°C with 20% O_2_. On day 2, NESC Maintenance medium was replaced with hMO Differentiation medium (DMEM/F-12 supplemented with 1× B-27 Supplement without Vitamin A, 1× N-2 Supplement, 1× GlutaMax, 1× Penicillin/Streptomycin, 10 ng/mL human BDNF (PeproTech, cat. No. 450-02), 10 ng/mL human GDNF (PeproTech, Cat. No. 450-10), 200 µM ascorbic acid (Sigma-Aldrich, Cat. No. A4544), 1 ng/mL TGF-β3 (Thermo Fisher Scientific Cat. No. 100-36E), 500 µM dibutyryl-cAMP (BioSynth, Cat. No. ND07996), and 1 µM PMA. Medium was changed again on day 4. On day 6, organoids designated for embedding were transferred into non-treated 24-well tissue culture plates (one organoid per well) using cut P1000 pipette tips to minimise mechanical damage. hMOs not intended for embedding were maintained in 96-well plates, and from day 8, the medium was switched to hMO Differentiation medium without PMA. For hMO embedding, 30 µL of undiluted Geltrex was added around each organoid, ensuring that organoids remained centered within droplets. Embedded hMOs were transferred into 24-well tissue culture plates (one organoid per well). DMEM/F-12 was replaced with 300 µL hMO Differentiation medium without PMA. From day 9 onward, embedded hMOs were cultured on an orbital shaker at 80rpm, while non-embedded hMOs were maintained under static conditions. Medium was changed every 3-4 days for both embedded and non-embedded hMOs until the end of culture.

### Senescence-associated β-galactosidase activity measurement

Staining was performed on astrocytes on day 24 of differentiation with the Senescence Detection Kit (Abcam, ab65351) following the manufacturer’s instructions. Images were acquired at 10X on an Olympus IX83 microscope and analysed with a custom MATLAB script.

### Immunohistochemistry of NESCs

Two-dimensional cultures of NESCs were washed once with PBS (Thermo Fisher, Cat. No. 10010023), fixed with 4% paraformaldehyde (PFA) (Millipore, Cat. No.1.00496.5000) for 15 minutes at room temperature, washed three times with PBS, and stored at 4°C in PBS until use.

Cells were then permeabilised for 15 minutes at RT in PBS containing 0.3% Triton X-100 (Carl Roth, Cat. No. 3051.3), washed 3 times with PBS and incubated for 1 hour at room temperature in blocking buffer (PBS containing 5% NGS (Abcam, Cat#ab138478)). Cells were then incubated with primary antibodies (antibodies list and concentrations in Material and Equipment Table) in 3% NGS in PBS blocking buffer overnight at 4°C. Cells were washed three times with PBS and incubated for 1 hour at room temperature in 3% NGS in 1x PBS with secondary antibodies (antibodies list and concentrations in Material and Equipment Table) and 1 µg/mL DAPI (Millipore, Cat#D8417). Finally, cells were washed three times with PBS and stored at 4°C in 1x PBS until imaging.

For the young (Y) condition, NESCs were at passage 5 (batch 1), passage 6 (batch 2), and passage 7 (batch 3). For the aged (A) condition, NESCs were analysed at passage 15 (batch 1), passage 16 (batch 2), and passage 17 (batch 3).

### Immunohistochemistry of astrocytes

Two-dimensional cultures of astrocytes were washed once with PBS (Thermo Fisher, Cat. No. 10010023), fixed with 4% paraformaldehyde (PFA) for 15 minutes at room temperature, washed three times with PBS, and stored at 4°C in PBS until use.

Cells were then permeabilised for 15 minutes at RT in 0.5% Triton X-100 in PBS for 15 minutes, washed once with 0.1% PBS-Tween (Applichem, Cat. No. A1389,0500) and incubated for 1 hour at room temperature in blocking buffer (PBS with 2% FBS and 0.1% Tween). Cells were then incubated with primary antibodies (antibody list and concentrations in Supplementary Table 1) in PBS with 2% FBS and 0.1% Tween blocking buffer overnight at 4°C. Cells were washed three times with PBS and incubated for 1 hour at room temperature in 2% FBS and 0.1% Tween in 1x PBS with secondary antibodies (antibodies list and concentrations in Material and Equipment Table) and Hoechst 33342 (Thermo Fisher, Cat#62249). Finally, cells were washed three times with 1x PBS and stored at 4°C in 1× PBS until imaging. For the young (Y) condition, astrocytes were generated from NSCs at passage 3 (batch 1), passage 4 (batch 2), and passage 5 (batch 3). For the aged (A) condition, astrocytes generated from NSCs at passage 11 (batch 1), passage 12 (batch 2), and passage 13 (batch 3) were analysed.

### Immunohistochemistry of hMOs

All immunohistochemical analyses of hMOs were performed on embedded hMOs. hMOs were washed once with PBS, fixed with 4% paraformaldehyde (PFA) overnight on an orbital shaker, washed three times with PBS, and stored at 4°C in 1× PBS containing 0.02% sodium azide (Carl Roth, Cat. No. K305.1) until use. Fixed hMOs were embedded in 3% low-melting-point agarose (Biozym, Cat. No. 840100) and sectioned at 80 µm using a vibratome. Sections were stored at 4°C in 1× PBS containing 0.02% sodium azide until use. For immunohistochemical analysis, free-floating two sections/immunostaining from 4-5 hMOs (lines HT, PD2, and PD1 GC) or 4-8 hMOs (line PD2) per batch were used for each antibody combination. Sections were permeabilised for 30 minutes at room temperature in PBS containing 0.5% Triton X-100. Sections were blocked for 2 hours at room temperature in 1PBS containing 5% NGS and 0.01% Triton X-100. Primary antibodies (antibodies list and concentrations in Supplementary Table 1) were diluted in 1× PBS containing 5% NGS and 0.01% Triton X-100, and sections were incubated for 48 hours at 4°C. Sections were then washed three times for 10 minutes each in 1× PBS containing 0.01% Triton X-100, then incubated with secondary antibodies (antibodies list and concentrations in Material and Equipment Table) and 1 µg/mL DAPI in PBS containing 5% NGS and 0.01% Triton X-100 for 2 hours at room temperature. Finally, sections were washed three times for 10 minutes each in PBS containing 0.01% Triton X-100, rinsed once with water, and mounted on microscopy slides with Fluoromount-G mounting medium (Thermo Fisher, Cat. No. 00-4958-02). All incubation and washing steps were performed with constant agitation on an orbital shaker.

For the young (Y) condition, hMOs were generated from NESCs at passage 5 (batch 1), passage 6 (batch 2), and passage 7 (batch 3). For the aged (A) condition, hMOs were generated from NESCs at passage 15 (batch 1), passage 16 (batch 2), and passage 17 (batch 3). hMOs at day 70 of differentiation were used for all the immunohistochemistry analysis.

### Image acquisition

For high-content imaging, live and fixed astrocytes cultured in 96-well plates were imaged with a Yokogawa CellVoyager CV8000 microscope (Yokogawa, RRID:SCR_023270). A 4x pre-scan allowed us to verify cell density, enabling the creation of a mask comprising 20 regions of interest (ROIs) per well to optimise acquisition. This mask guided the device during high-resolution acquisition with a 63x objective. hMOs sections were also acquired using a Yokogawa CellVoyager CV8000 microscope with a 20x objective.

NESC immunostaining photos were acquired manually using a 20x or a 63x (only for TOM20 immunostaining) objective, selecting five random ROIs for each well. For all the NESCs, immunostaining an automated inverted Nikon Ti-E microscope, except for TOM20, for which the Leica TCS SP8 STED microscope was used.

### Analysis of NESC immunostaining

Multichannel fluorescence microscopy data were analysed using a custom Python pipeline (code available on GitHub: link here). Data was batch-processed as multichannel z-stacks in CZYX format. For each dataset, all available stacks were processed iteratively, and the images within each stack were analysed sequentially. Individual channels were extracted, z-projections were generated, and images were processed through marker-specific quantification workflows. Output included quantitative measurements in tabular format and quality control visualisations for validation of segmentation and measurements. Pairwise comparisons between young (Y) and aged (A) samples within each cell line were performed using two-sided Mann–Whitney U tests in R with the wilcox.test() function from the stats package (exact = FALSE).

### Analysis of astrocytes and hMOs immunostaining

For quantitative image analysis of astrocytes as well as for hMO, each acquired photo was analysed using a custom image-analysis algorithm in MATLAB (2021, Mathworks, RRID:SCR_001622). Pairwise comparisons between young (Y) and aged (A) samples within each cell line were performed using two-sided Mann–Whitney U tests in R with the wilcox.test() function from the stats package (exact = FALSE).

### Representative images processing

Representative immunofluorescence images in Fig. 1b, h, m, o and Extended Fig. 1c, e were processed using Fiji/ImageJ. ND2 image files were imported using the Bio-Formats plugin and visualized as composite images. For each sample, a representative z-plane was selected and exported for presentation, except for Extended Data Fig. 1e, for which a maximum-intensity projection was generated. Channel-specific contrast was initially adjusted on a reference image, and the resulting brightness and contrast settings were subsequently applied identically to all images within the dataset. Representative immunofluorescence images shown in Fig. 2b, f, h, j,l, n, r and Fig. 3b, h, j were exported from the MATLAB-based image analysis pipeline used for immunocytochemistry (ICC) quantification.

**Figure 1:**
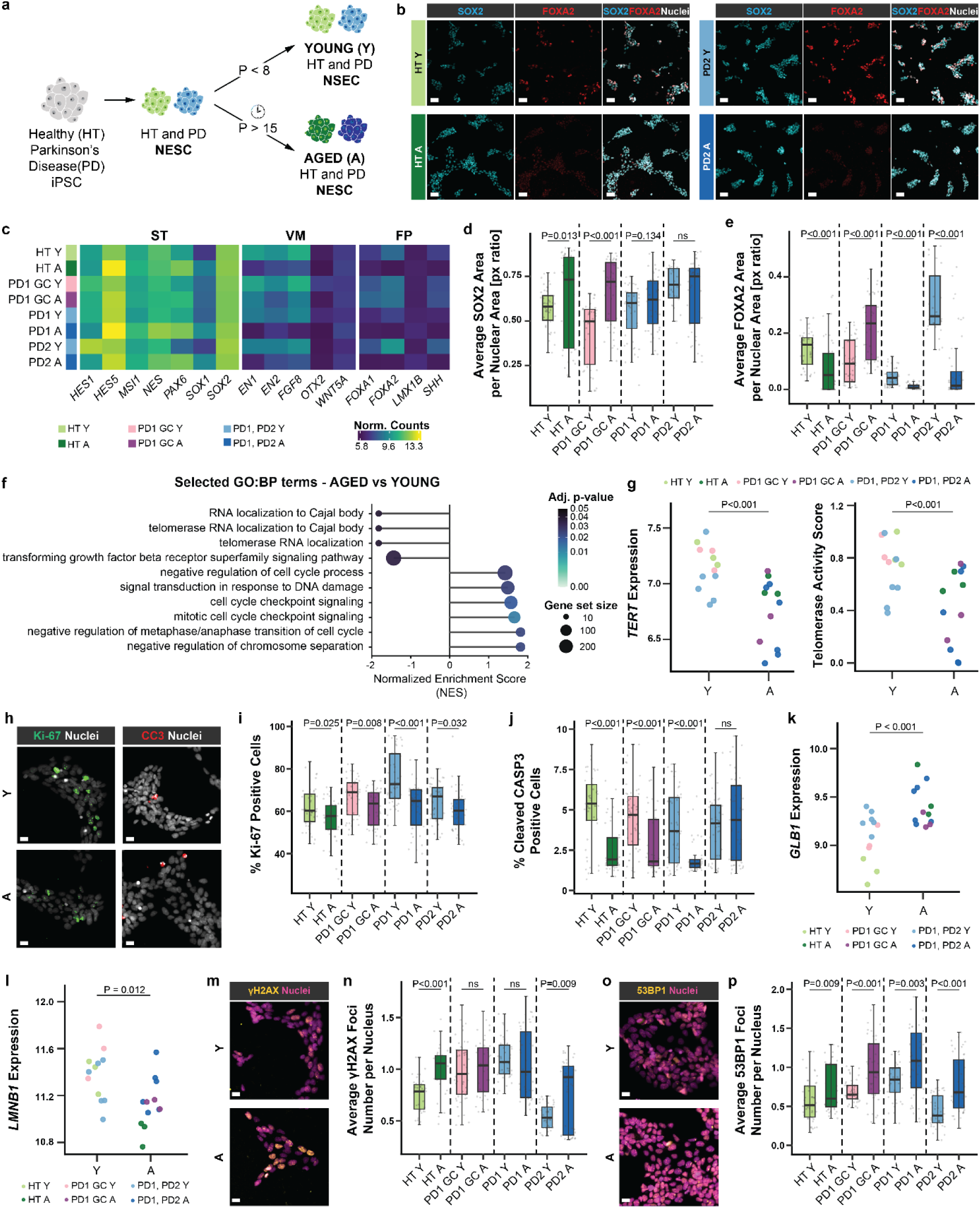
Extensive passaging induces line-dependent senescence-like features in NESCs while preserving lineage identity. (**a**) Schematic overview of the passage-induced replicative exhaustion approach applied to human iPSC-derived NESCs, where P denotes passage numbers. (**b**) Representative immunofluorescence images of NESC identity markers. Single channels: SOX2 (red) and FOXA2 (cyan). Merged image includes SOX2 (red), FOXA2 (cyan), and Nuclei (DAPI, grey). Scale bars, 140 μm. (**c**) mRNA expression of canonical ventral NESC markers. ST, stemness markers; VM, ventral midbrain patterning markers; FP, floor plate progenitor markers. Expression values shown as variance-stabilising transformed (VST)-normalised counts. (**d**, **e**) Quantification of SOX2 (**d**) and FOXA2 (**e**) mean positive area normalised per nuclear area. (**f**) Lollipop plot showing selected Gene Ontology Biological Process (GO-BP) terms enriched in differentially expressed genes between Y and A NESCs using gene set enrichment analysis (GSEA). Circle size indicates gene set size; colour indicates adjusted P-value; x-axis shows normalised enrichment score (NES). (**g**) Left: VST-normalised counts of *TERT* in Y and A conditions. Right: telomerase activity score calculated using the EXTEND R package ^33^. (**h**) Representative immunofluorescence images of Ki-67 (left) and cleaved caspase-3 (CC3, right) in Y (top) and A (bottom) HT NESCs. Merged images show Ki-67 (green) with DAPI (grey) (left set) and CC3 (red) with DAPI (grey). Scale bars, 20 μm. (**i**, **j**) Quantification of the percentage of Ki-67-positive (**i**) and cleaved caspase-3-positive (**j**) NESCs. Percentages for each data point are expressed as the number of positive nuclei over the total number of nuclei. (**k, l**) VST-normalised counts of *GLB1* (**k**) and *LMNB1* (**l**) expression in Y and A conditions. (**m**) Representative immunofluorescence images of γH2AX nuclear foci in Y (top) and A (bottom) HT NESCs. Merged images show γH2AX (yellow) with DAPI (magenta). Scale bars, 20 μm. (**n**) Quantification of mean γH2AX foci number per nucleus. (**o**) Representative immunofluorescence images of 53BP1 nuclear foci in Y (top) and A (bottom) HT NESCs. Merged images show 53BP1 (yellow) with DAPI (magenta). Scale bars, 20 μm. Representative images were contrast-adjusted separately for visualisation. (**p**) Quantification of mean 53BP1 foci number per nucleus. For immunofluorescence-based analyses, three independent NESC batches per condition were analysed (Y: passages 5-7; A: passages 19-21). Each dot represents an individual image acquisition. For RNA sequencing, three independent NESC batches per condition were analysed (Y: passages 5-7; A: passages 17-19), and each dot represents one sequenced sample. HT, healthy control; PD1 and PD2, Parkinson’s disease lines 1 and 2; PD1 GC, gene-corrected isogenic control of PD1 line; Y, young cells (<8 passages); A, aged cells (>15 passages). Mann-Whitney U tests were used for pairwise comparisons between Y and A within each genetic background in (d), (e), (i), (j), (n), and (p). Differential gene expression analysis for (g), (k), and (l) was performed using the DESeq2 R package. Telomerase activity score in (g) was analysed using a linear mixed-effects model.

**Figure 2:**
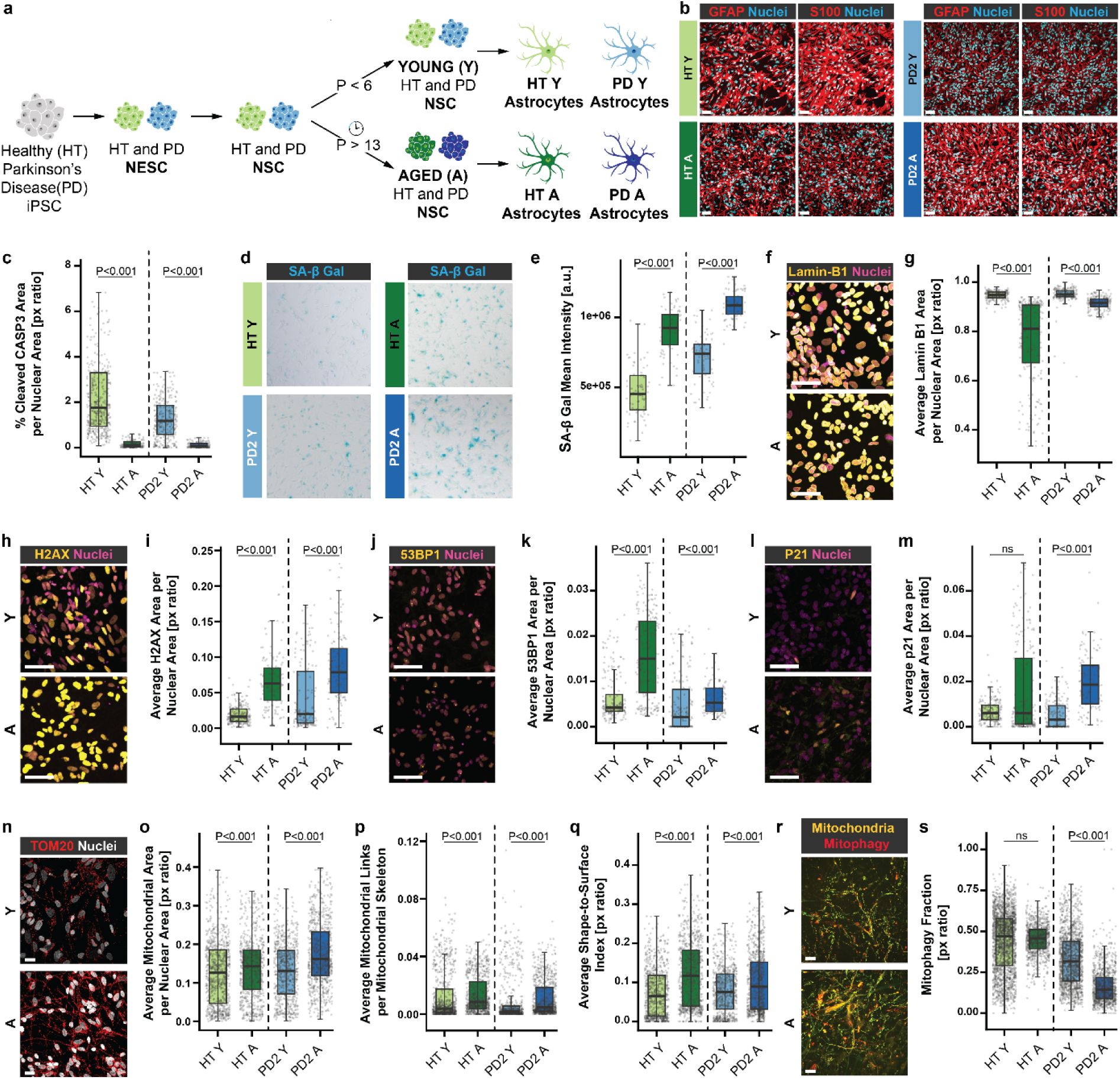
Extensive passaging of NSCs induces line-dependent senescence-like features in astrocytes while preserving cellular identity. (**a**) Schematic overview of the passage-induced replicative exhaustion approach applied to human iPSC-derived neural stem cells (NSCs) prior to astrocyte differentiation, where P denotes passage number. (**b**) Representative immunofluorescence images of astrocyte identity markers. Merged images show GFAP (red, left panels of each set) with Nuclei (DAPI, cyan) and S100B (red, right panels of each set) with Nuclei (DAPI, cyan). Scale bars, 60 μm. (**c**) Quantification of the percentage of cleaved caspase-3-positive area. Percentages for each data point are expressed as the number of positive nuclei over the total number of nuclei (**d**). Representative images of senescence-associated β-galactosidase (SA-β-gal, blue deposits) activity in Y and A astrocytes. Scale bars, xx μm. (**e**) Quantification of SA-β-gal activity as arbitrary intensity units (a.u.). (**f**) Representative immunofluorescence images of Lamin B1 in Y (top) and A (bottom) HT astrocytes. Scale bars, 60 μm. (**g**) Quantification of mean Lamin B1 fluorescence intensity per nucleus. (**h**) Representative immunofluorescence images of γH2AX nuclear foci in Y (top) and A (bottom) HT astrocytes. Scale bars, 60 μm. (**i**) Quantification of H2AX area relative to the total nuclear area. (**j**) Representative immunofluorescence images of 53BP1 nuclear foci in Y (top) and A (bottom) HT astrocytes. Scale bars, xx μm. (**k**) Quantification of 53BP1 foci area relative to the total nuclear area. (**l**) Representative immunofluorescence images of p21 in Y (top) and A (bottom) HT astrocytes. Scale bars, 60 μm. (**m**) Quantification of p21-positive area relative to the total nuclear area. (**n**) Representative immunofluorescence images of the outer mitochondrial membrane marker TOM20 in Y (top) and A (bottom) HT astrocytes. Scale bars, 20 μm. (**o**) Quantification of mitochondrial area normalised to the total nuclear area. (**p**) Mitochondrial branching quantified as the number of skeleton links normalised to mitochondrial skeleton. (**q**) Mitochondrial elongation quantified as the shape-to-surface ratio derived from erosion-separated mitochondrial bodies. (**r**) Representative images of the mitophagy reporter in Y (top) and A (bottom) HT astrocytes. Scale bars, 20 μm. (**s**) Quantification of the mitophagy fraction expressed as a ratio between the mitophagy signal and the total mitochondria network signal. For immunofluorescence-based analyses, three independent astrocyte batches per condition were analysed (Y: young astrocytes derived from NSC passages 3-5; A: aged astrocytes derived from NSC passages 14-16). Each dot represents one independent image acquisition. HT, healthy control; PD2, Parkinson’s disease line 2. Mann-Whitney U tests were used for pairwise comparisons between Y and A within each cell line in (c), (e), (g), (i), (k), (m), (o), (p), (q) and (s).

**Figure 3:**
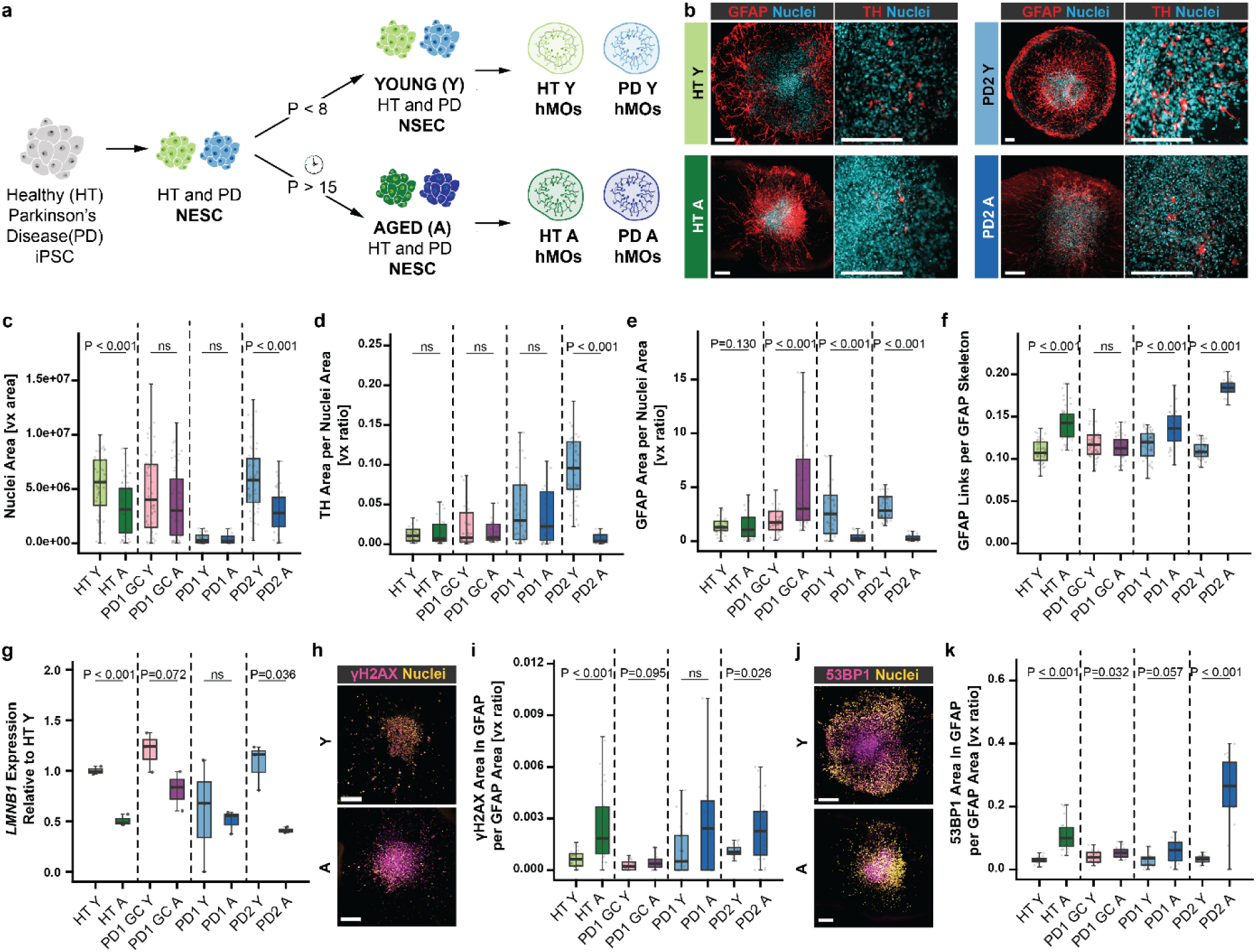
Extensive passaging of NESCs induces line-dependent senescence-like features in hMOs. (**a**) Schematic overview of the passage-induced replicative exhaustion approach applied to NESCs, where P denotes passage numbers, prior to hMOs generation. (**b**) Representative immunofluorescence images of astrocyte and dopaminergic neuron markers in hMOs. Merged images show GFAP (red) with DAPI (cyan) (left set) and TH (red) with DAPI (cyan) (right set). Scale bars, 200 μm. (**c**) Quantification of mean nuclear area. (**d**) Quantification of TH-positive area relative to the total nuclear area. (**e**) Quantification of GFAP-positive area relative to the total nuclear area. (**f**) Quantification of astrocyte branching, expressed as GFAP skeleton link number relative to the total GFAP-positive area. (**g**) *LMNB1* mRNA expression normalized to *RPL37A* and expressed relative to HT Y. (**h**) Representative immunofluorescence images of γH2AX nuclear foci in Y (top) and A (bottom) HT hMOs. Scale bars, 200 μm. (**i**) Quantification of γH2AX foci total area within GFAP-positive regions in immunofluorescence images relative to the total GFAP area. (**j**) Representative immunofluorescence images of 53BP1 nuclear foci in Y (top) and A (bottom) HT hMOs. Scale bars, 200 μm. (**k**) Quantification of 53BP1 foci total area within GFAP-positive regions relative to the total GFAP area. For immunofluorescence-based analyses, three independent hMO batches per condition were analysed (Y: hMOs derived from NESCs passages 5-7; A: hMOs derived from NESCs passages 17-19). Each dot represents one independent image of an hMO section. For each batch, sections from at least four independent hMOs were analysed (two sections per organoid). For qPCR analysis, three independent hMO batches per condition were analysed (Y: passages 5-7; A: passages 17-19), each comprising pooled samples from at least 8 organoids. Mann-Whitney U tests were used for pairwise Y versus A comparisons within each genetic background in (d), (e), (f), (j), and (l). Two-sided t-tests were used for mRNA analyses in (g) and (h). HT, healthy control; PD1 and PD2, Parkinson’s disease lines; PD1 GC, gene-corrected isogenic control of PD1.

### RNA extraction from hNESCs and hMOs

NESCs pellets (250000-500000 per individual sample) or non-embedded hMOs at day 70 of differentiation (5-8 per individual sample) were washed in PBS, snap frozen, and stored at -80°C until used. For the RNA extraction of the hMOs, first 1mL of TRIzol (Thermo Fisher, Cat. No. PR94757) was added per sample, the pellet was thoroughly lysed by pipetting, and the lysate was incubated for 5 minutes at room temperature. Following, the lysate was transferred to a Phasemaker tube (Thermo Fisher, Cat. No. 15645268), 200 µl of Chloroform was added, emulsified by vigorous inversion and incubated for 5 minutes at room temperature. Solutions were then centrifuged at 4°C with a speed of 12000x*g* for 15 minutes. Then the aqueous phase containing the RNA is transferred and mixed with one volume of 70% Ethanol. Afterwards, the resulting solution was loaded onto Qiagen RNeasy kit columns (Qiagen, Cat No. 74106), and the RNA purification was performed according to the manufacturer’s instructions. The RNA was eluted in nuclease-free water in the final step. All RNA samples extracted this way were treated with DNase I (Sigma Aldrich, Cat. No. AMPD1) according to manufacturer instructions before proceeding with the cDNA generation.

Alternatively, for the NESCs or the non-embedded hMOs RNA extraction, the Maxwell^®^ RSC simplyRNA Tissue Kit (Promega, Cat. No. AS1340) was also used according to manufacturer instructions, for which the DNase I treatment is already included in the procedure, so no extra treatment was performed. RNA was eluted in nuclease-free water.

The RNA quantity and quality were evaluated through NanoDrop analysis, and it was stored at -80°C to be used for the following steps.

### cDNA generation and qPCR

RNA conversion to cDNA was performed according to manufacturer instructions using (Bio-Rad, Cat. No. 1708890) or (Thermo Fisher, Cat. No. 4387406) and stored at -20°C until use. All qPCR reactions were performed using a total amount of 3-15ng of RNA in a final volume of 10 µl per reaction with a total of 3 technical replicates per reaction. qPCR was performed by using TaqMan Gene Expression Assay (FAM) (refer to the Material and Equipment Table for the complete list of primer pairs used) together with TaqMan^™^ Universal PCR Master Mix II, no UNG (Applied Biosystems, Cat. No. 4440040) according to the manufacturer’s instructions. Alternatively, qPCR was performed by using custom primer pairs (refer to the Material and Equipment Table for the complete list of primer pairs used) together with iQ SYBR^®^ Green Supermix (Bio-Rad, Cat. No. 1708880). qPCR experiments were run on a Bio-Rad CFX384 Real-Time PCR Detection System. For each gene-sample combination, three technical replicates were made, and the mean Ct value was used for downstream analysis. Gene-sample combinations with fewer than two technical replicates yielding detectable Ct values were excluded from the final analysis. Relative gene expression was quantified using the ΔΔCt method, with *RPL37A* serving as the housekeeping gene for ΔCt normalisation, while the mean ΔCt value of the HT Y group was used as the calibrator for ΔΔCt calculations and for generating the relative expression values shown in the plot. Statistical analyses were conducted on the normalised values in R using the Student’s t-test (stats package; t.test (group1, group2, var.equal = TRUE, paired = FALSE)).

For the young (Y) condition, hMOs were generated from NESCs at passage 5 (batch 1), passage 6 (batch 2), and passage 7 (batch 3). For the aged (A) condition, two independent hMOs derivation were generated from NESCs at passage 15 (batch 1), passage 16 (batch 2), and passage 17 (batch 3) and at passage 17 (batch 1), passage 18 (batch 2), and passage 19 (batch 3). Passages are here considered biological replicates.

### hMOs lipidomic

Lipids and proteins were extracted using a methyl tert-butyl ether (MTBE)-based method adapted from^31^.

Briefly, 5-8 non-embedded hMOs were collected per sample in 50 µl of 50 mM ammonium bicarbonate (ABC). Cells were lysed by three freeze-thaw cycles (freezing at -80 °C and thawing in an ultrasonic bath). Methanol (MeOH) and MTBE were added, and samples were incubated in a thermomixer (Grant Instruments) at 500 rpm for 60 minutes. Phase separation was induced by the addition of water to obtain a final MTBE:MeOH:H₂O ratio of 3.3:1:3 (v/v/v), followed by centrifugation at 1,000xg for 10 minutes. The upper organic phase containing lipids was collected. The lower phase was re-extracted with the solvent corresponding to the composition of the upper phase. After centrifugation, the upper phase was collected and combined with the initial organic extract. The combined organic phases (lipid fraction) and the remaining lower phase, including the pellet (protein fraction), were dried in a vacuum centrifuge. The protein fraction was reconstituted in 5 M urea in 50 mM ABC. Insoluble material was removed by centrifugation at 30,000xg for 15 minutes, and the supernatant was used for Bradford assay-based protein quantification. The lipid fraction was normalised to protein content, reconstituted in isopropanol/acetonitrile (IPA/ACN, 50:50, v/v), and stored at -20 °C until LC-MS analysis.

LC-MS lipidomics analysis was adapted from ^32^. Lipids were separated using an Ultimate 3000 Rapid Separation UHPLC system (Thermo Scientific Dionex) equipped with a Hypersil Gold C18 analytical column (10 cm × 2.1 mm, 1.9 µm; Thermo Scientific). Separation was achieved using an 18-minute gradient from 68% mobile phase A (ACN:H₂O 60:40 (v/v) containing 10 mM ammonium formate (AF)) to 97% mobile phase B (IPA:ACN 90:10 (v/v) containing 10 mM AF) at a flow rate of 0.25 ml min⁻¹. The UHPLC system was coupled to a Q Exactive HF mass spectrometer (Thermo Scientific). Data were acquired in data-dependent acquisition (DDA) mode. Full MS scans were recorded over a mass range of m/z 200-1,450 at a resolution of 60,000, followed by MS/MS acquisition of the top eight most intense ions at a resolution of 30,000 with dynamic exclusion set to 60 seconds. Data analysis was performed using Lipostar (v2.1.9).

For the young (Y) condition, hMOs were generated from NESCs at passage 5 (batch 1), passage 6 (batch 2), and passage 7 (batch 3). For the aged (A) condition, hMOs were generated from NESCs at passage 17 (batch 1), passage 18 (batch 2), and passage 19 (batch 3). Passages are here considered biological replicates.

### hMOs lipidomic data analysis

Raw lipid intensity data exported from Lipostar were imported into R. Lipids were retained for downstream analysis if non-zero intensity values were present in at least 35% of samples across the dataset. To characterise overall shifts in lipid class abundance (Extended figure 4b), raw intensities were summed per lipid category and per sample, and category totals were log2-transformed before statistical testing. Results were visualised as a stacked bar plot of proportional log2 category totals per sample, with individual batch-level values overlaid as points.

Differential abundance analysis of individual lipids was performed using the limma R package. A design matrix encoding cell line (HT, PD1 GC, PD1, and PD2) and treatment (YOUNG and AGED) was constructed, and a linear model was fitted to the log2-transformed intensity matrix using lmFit(). Contrast matrices encoding four AGED-YOUNG comparisons, one per line (HT, PD1 GC, PD1, and PD2), were defined and fitted using contrasts.fit(), followed by empirical Bayes moderation via eBayes(). To summarise differential abundance at the lipid subclass level, limma t-statistics were averaged across all lipids within each subclass, and the resulting subclass-level effect estimates across all four contrasts were displayed as a heatmap using the ComplexHeatmap R package, with a diverging colour scale and BH-adjusted significance annotations.

Lipid ontology over-representation analysis (ORA) was performed using the LION/web tool. For HT, PD1 GC, and P1 and PD2 together, a ranked list of lipids was derived from the corresponding limma differential abundance results (AGED-YOUNG contrast). Lipid intensities were first aggregated at the lipid ID level by summing duplicate measurements per sample, log₂-transformed after addition of a pseudocount (+1) and organised into a sample-by-lipid matrix. Linear modelling was performed using the limma framework with empirical Bayes moderation. For each contrast, lipids were ranked using a signed statistic defined as the product of the adjusted p-value and the sign of the log₂ fold change (sign(logFC) × adj.P.Val). Enriched LION terms were identified at a BH-adjusted p-value threshold of 0.05. For each genotype, the four terms with the lowest FDR were selected for visualisation. Enrichment results were displayed as a dot plot in which the x-axis encodes −log10(FDR), point size encodes the number of annotated lipids overlapping the term, and panels are faceted by HT, PD1 GC and PD groups. Weighted gene co-expression network analysis (WGCNA) was applied to the filtered, log2-transformed lipid intensity matrix to identify co-regulated lipid modules. The network was constructed using blockwiseModules() from the WGCNA R package with a signed topological overlap matrix (TOM), biweight midcorrelation (bicor) as the similarity measure, a soft-thresholding power of 6, a minimum module size of 10 lipids, deepSplit = 4, and a merge cut height of 0.25. To quantify the association of each module with aging and cell line identity, a trait matrix was constructed comprising a binary ageing variable (YOUNG = 0, AGED = 1) and one-hot-encoded cell line indicators. For each module, a linear model, lm(ME ∼ CellLine * Ageing) was fitted on the module eigengene values, from which the ageing β coefficient and the Line × Ageing interaction F-statistic were extracted. Multiple testing correction across modules was applied independently for the ageing and interaction p-values using the Benjamini-Hochberg method. Module eigengene values were visualised as a heatmap (modules as rows, samples as columns) using the ComplexHeatmap R package and (Extended Figure 4c). Module-level ageing associations were displayed as a two-panel heatmap (Figure 4b) in which one column encodes the ageing β coefficient per module and a second column encodes the −log10 BH-adjusted p-value of the Line × Ageing interaction. Hub lipids within the two most ageing-associated modules were identified by module membership (kME), defined as the Pearson correlation between each lipid’s expression profile and the module eigengene (Figure 4c). The lipid class composition of each module was displayed as a stacked proportional bar chart, and the top 10 hub lipids ranked by kME in each module were displayed as horizontal bar charts with log2 fold change annotations and BH-adjusted significance overlaid.

**Figure 4:**
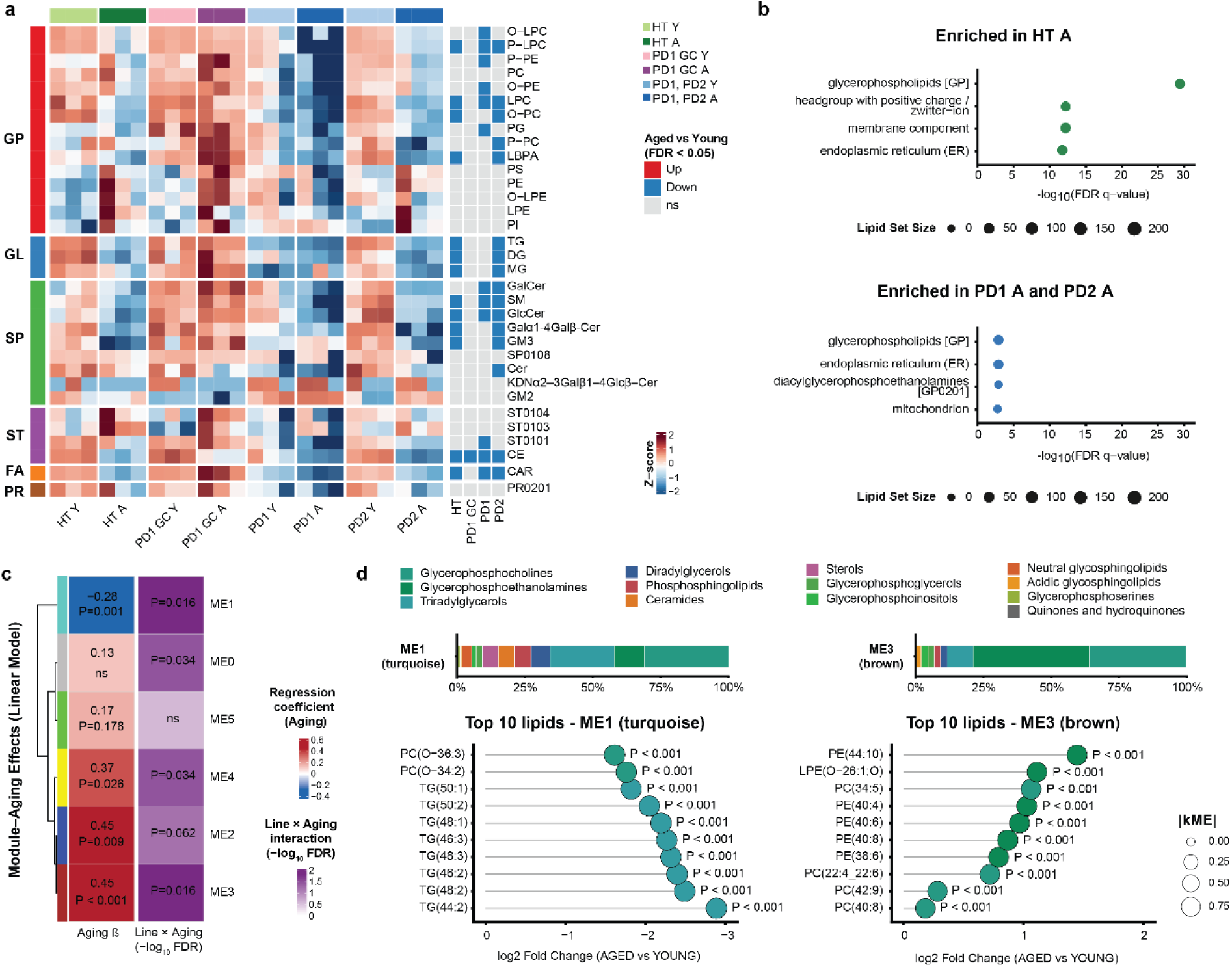
Extensive passaging of NESCs induces line-dependent lipidome changes in hMOs. (**a**) Heatmap of lipid class abundances across hMO samples. Abundances were log10(x+1) transformed and Z-score scaled per lipid subclass. Blue, red, and grey squares indicate differential abundance in pairwise Y versus A comparisons within each genetic background. Lipid classes: GP, glycerophospholipids; GL, glycerolipids; SP, sphingolipids; ST, sterols; FA, fatty acyls; PR, prenol lipids. (**b**) Lipid Ontology (LION) enrichment analysis of differentially abundant lipid species in Y versus A comparisons for HT hMOs (top) and PD1 + PD2 combined hMOs (bottom). X-axis indicates log10(FDR q-value); dot size indicates number of lipids per term. (**c**) Heatmap summarising module-level linear model results derived from WGCNA module eigengenes (MEs). Linear models were fitted as ME ∼ Line × Aging. The right column displays the estimated β coefficient for the Aging term (adjusted for Line, top value), indicating the direction and magnitude of ageing-associated changes per module. Values are annotated with Benjamini-Hochberg-adjusted P values (bottom values). The left column shows the -log10(FDR) of the Line × Aging interaction term (ANOVA F-test), reflecting genotype-dependent differences in the aging effect. Significant interaction terms indicate that the magnitude of ageing-related changes differs between cell lines. (**d**) Module composition and 10 top lipid drivers for representative modules (ME1 and ME3). Upper panels show stacked bar plots of lipid class composition within each module. Lower panels show lollipop plots of 10 top lipid species ranked by module eigengene-based connectivity (kME, correlation with the module eigengene). Lollipop length represents log2 fold change (A vs Y); dot size indicates |kME|; dot colour indicates lipid class; adjusted P values from differential abundance analysis are indicated. Three independent hMOs batches per condition were analysed (Y: passages 5-7; A: passages 17-19), each comprising pooled samples from 5-8 organoids Differential lipid abundance was assessed using linear modelling with Benjamini-Hochberg correction. HT, healthy control; PD1 and PD2, Parkinson’s disease lines; PD1 GC, gene-corrected isogenic control of PD1.

### NESCs bulk RNA-sequencing data generation

For the young (Y) condition, NESCs were analysed at passage 5 (batch 1), passage 6 (batch 2), and passage 7 (batch 3). For the aged (A) condition, NESCs were analysed at passage 17 (batch 1), passage 18 (batch 2), and passage 19 (batch 3).

Frozen RNA samples were shipped to Singleron Technologies GmbH (Cologne, Germany) for transcriptome sequencing. RNA quantity and integrity were assessed using the Qubit RNA High Sensitivity (HS) Assay Kit (Invitrogen, Cat. No. Q10211) and the High Sensitivity RNA ScreenTape system (Agilent Technologies, Cat. No. 5067-5579) prior to library preparation using the AccuraCode RNA-Seq Kit V2 (Singleron Biotechnologies GmbH, Cat. No. 10710174) according to the manufacturer’s instructions.

Briefly, polyadenylated mRNA was captured using paramagnetic beads. During reverse transcription, 100 ng mRNA was converted to cDNA and labelled with sample-specific barcodes and unique molecular identifiers. Barcoded cDNA from all samples was pooled and amplified. A total of 50 ng cDNA was used for library construction. Libraries were sequenced on a NovaSeq X platform (Illumina) using paired-end 150-bp sequencing by GENEWIZ (Leipzig, Germany).

Raw sequencing data were processed into gene expression matrices using CeleScope v2.7.4 (multi_bulk_rna workflow with customised chemistry; https://github.com/singleron-RD/CeleScope). R2 FASTQ files containing transcript sequences were demultiplexed using well-specific barcodes extracted from the corresponding R1 FASTQ files. Reads were aligned to the *Homo sapiens* reference genome (GRCh38, Ensembl release 99) using STAR. Bulk RNA sequencing data generated in this study have been deposited in the Gene Expression Omnibus (GEO) database under accession number GSE331395.

### NESCs bulk RNA-sequencing data analysis

Gene count matrices were imported into R, and a DESeq2 dataset was constructed using DESeqDataSetFromMatrix() with the design formula ∼ CellLine + Ageing + Batch to account for cell line identity (i.e. HT, PD1 GC, PD1, PD2) and experimental batch. Low-count genes were removed by retaining only genes with a minimum of ten counts in at least three samples. Variance-stabilising transformation (VST) was applied to all retained genes using vst() from the DESeq2 package for visualisation (Fig. 1c-j-k-l).

Differential expression analysis was performed using DESeq() with the design formula described above. Genes with a Benjamini-Hochberg adjusted p-value below 0.05 and an absolute log2 fold change greater than 1 were classified as differentially expressed. The Wald test statistic from the DESeq2 output was used as the gene-level ranking metric for gene set enrichment analysis. Gene Set Enrichment Analysis (GSEA) was performed on the ranked gene list using gseGO() from the clusterProfiler R package, targeting Gene Ontology Biological Process (GO:BP) terms annotated for Homo sapiens via the org.Hs.eg.db annotation database. A curated subset of biologically relevant terms was selected for visualisation and displayed as a lollipop plot in which the Normalised Enrichment Score (NES) is shown on the x-axis, point size encodes gene set size, and point colour encodes the BH-adjusted p-value on a square-root-transformed scale (Fig. 1f).

VST-normalised expression levels of individual genes of interest (*TERT*, *LMNB1* and *GLB1*) were visualised as jitter plots grouped by ageing conditions independent of the cell line (YOUNG vs AGED). Statistical significance between the YOUNG and AGED groups was reported as computed and adjusted by the DESeq2 package.

Telomerase activity was estimated from VST-normalised expression data using the EXTEND package ^33^ (Fig. 1g, right). Statistical significance of the YOUNG vs AGED difference was assessed using a linear mixed model lmer(NormEXTENDScores ∼ Ageing + (1|CellLine) + (1|Batch)) from the lme4 R package.

Gene sets enrichment (Extended Fig. 1b, Fig. 5a, and Extended Fig. 5a, b, c) were quantified using the single-sample Gene Set Enrichment Analysis (ssGSEA) R package ^34^ (Extended Fig. 1b). Gene sets were either manually curated based on published literature: Stem Identity (*SOX2, PAX6, HES5, ASCL1, SOX1, PAX3, DACH1, LMO3, NR2F1, PLAGL1, LIX1, HOXA2, FOXA2, IRX3, HES1, MSI1*), Ventral Midbrain identity (*SHH*, *FOXA1*, *FOXA2*, *GLI1*, *GLI2*, *PTCH1*, *NKX2-2*, *NKX6-1*, *NKX6-2*, *OTX2*, *LMX1A*, *LMX1B*, *EN1*, *EN2*, *WNT1*, *FGF8*, *CORIN*, *MSX1*, *NR4A2*, *PITX3, WNT5A*, *PBX1*), SASP (*IL6, CXCL8, IL1A, IL1B, CXCL1, CXCL2, CCL2, CCL20, MMP1, MMP3, MMP10, SERPINE1, IGFBP3, IGFBP7, TGFB1, VEGFA, CSF2*); or chosen from Gene Ontology (GO) database: chromatin organization (GO:0006325), cellular senescence (GO:0090398), cellular response to DNA damage stimulus (GO:0006974), mitochondrion organization (GO:0007005), response to oxidative stress (GO:0006979), mitophagy (GO:0000422), macroautophagy (GO:0016236), and lipid metabolic process (GO:0006629). Differences in pathway activity between YOUNG and AGED samples shown in Extended Fig. 1b were assessed using linear models fitted to ssGSEA enrichment scores. Pairwise contrasts were performed to compare AGED and YOUNG samples within each genotype group (HT, PD1 GC, PD1, and PD2). P values were adjusted for multiple testing using the Benjamini-Hochberg method, and pathways with a false discovery rate (FDR) < 0.05 were considered statistically significant.

**Figure 5:**
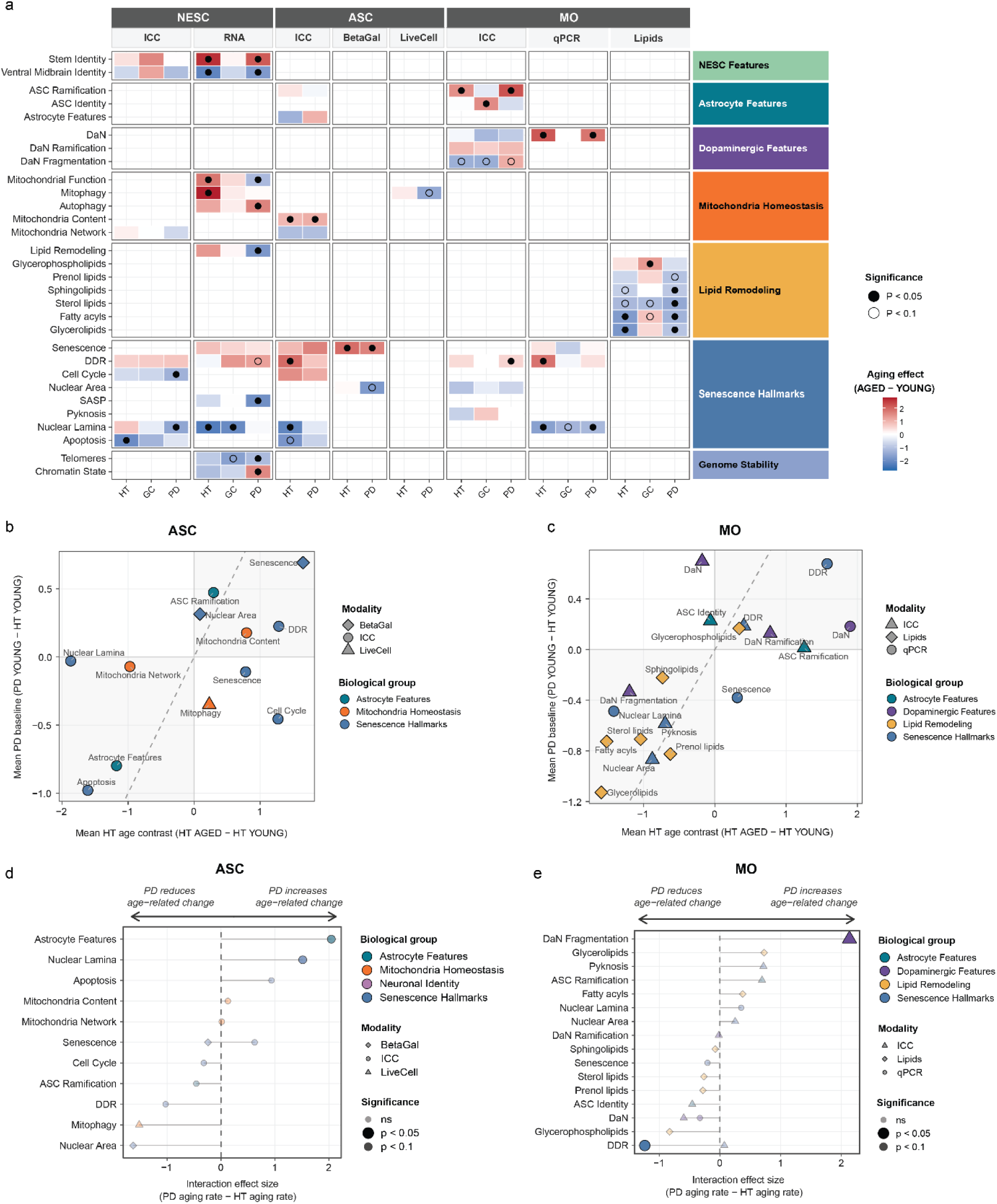
Category-level summary of aging and PD-associated effects in astrocytes and hMOs. (**a**) Heatmap of mean category scores (averaged z-scored features within each biological category × modality group) across cell lines (HT, PD, GC, with PD1 and PD2 cell lines analysed together) and passage conditions. Rows represent biological category × modality combinations; columns represent condition means. Colour scale indicates mean z-score relative to the HT Y reference per category. (**b, c**) Scatter plots of mean PD baseline effect (PD Y− HT Y; y-axis) versus mean HT aging effect (HT A − HT Y; x-axis) at the biological category level, for astrocytes (**b**) and hMOs (**c**). Each point represents one biological category; colour indicates biological group, and point shape indicates measurement modality. The grey regions highlight features that show concordant behaviour in PD Y and HT A relative to the HT Y baseline. The dashed diagonal represents equal effects of aging and PD relative to baseline. The dashed diagonal represents equal aging and PD baseline effects. (**d, e**) Lollipop plots showing the interaction effect between genotype (PD vs HT) and age on biological category scores in astrocytes (**d**) and hMOs (**e**). The interaction effect quantifies the degree to which PD amplifies or attenuates age-related change (PD aging rate - HT aging rate). Each point represents one biological category × modality; colour indicates biological group and shape indicates measurement modality. Point transparency and size reflect statistical significance (opaque/large: p < 0.05; intermediate: p < 0.1; faded/small: not significant). Horizontal lines connect each category to the zero reference. Statistical significance was assessed using linear mixed-effects models with genotype × age interaction terms. HT, healthy; PD, Parkinson’s disease; GC, gene-corrected of PD1; Y, young (low passage); A, aged (high passage); ICC, immunocytochemistry-derived morphometric features; qPCR, mRNA expression; Lipids, lipidomic features.

### Integrative multi-modal analysis

To characterise age-related changes across genotypes and cell types in an integrative framework, per-assay measurement tables from immunocytochemistry (ICC), bulk RNA-seq (RNA), quantitative PCR (qPCR), live cell (LiveCell), SA-β-gal (BetaGal), and lipidomics (Lipids) readouts were combined into unified long-format datasets for each cell type analysed in this study (NESCs, astrocytes, and hMOs). z-scoring was performed independently within each Modality for each feature using scale(), standardising all numeric features to zero mean and unit variance at the assay level.

Each feature was assigned to a biological category (e.g., DDR, Nuclear Lamina, Mitochondria Network, Stem Identity, Lipid Remodelling) and an overarching biological macro-group (e.g., Senescence Hallmarks, Mitochondria Homeostasis, Genome Stability). Category-level summary scores were derived by averaging z-scores across all features within a given category and modality.

To assess the cross-line consistency of ageing effects within the PD genotype, which comprised two independent lines (PD1 and PD2), a parallel model lm(Score_z ∼ CellLine * Age) was fitted per feature, with per-line ageing contrasts extracted via emmeans. An integrated cross-cell-type summary was produced as a heatmap combining NESCs, astrocytes, and hMOs. In this heatmap, each tile encodes the per-genotype ageing effect (AGED - YOUNG) at the category level, estimated by fitting a linear model on category-level mean z-scores and extracting per-genotype contrasts using emmeans.

To quantify the effect of aging and genotype on each feature and biological category, a linear model was fitted per feature-modality combination using lm(Score_z ∼ Genotype * Aging), where Genotype is HT, PD1 GC and PD1 and PD2 together. Estimated marginal means were computed using emmeans() from the emmeans package, from which three pre-specified planned contrasts were extracted: (i) the healthy aging contrast (HT AGED - HT YOUNG), quantifying the direction and magnitude of age-related change in healthy controls; (ii) the PD baseline shift (PD YOUNG - HT YOUNG), capturing steady-state differences between the PD-mutant and healthy genotypes independent of aging; and (iii) the PD × aging interaction ((PD AGED - PD YOUNG) - (HT AGED - HT YOUNG)), reflecting whether the PD mutation amplifies or attenuates the aging trajectory relative to healthy controls.

The gene-corrected (GC) genotype was included in both models to allow unbiased estimation of all marginal means, with a GC-specific ageing contrast extracted for the combined heatmap, where it serves as a visual validation layer. Nominal p-values were used for all main-figure significance overlays.

## Results

### Prolonged passaging induces stem cell exhaustion and senescence-like features in NESCs

To investigate whether passaging-induced exhaustion can model senescence-associated features *in vitro*, NESCs were analysed at low and high passages (Fig. 1a). Our experimental data set consisted of NESCs derived from a healthy individual (HT), and two PD patients carrying the LRRK2 G2019S mutation (PD1, PD2), and an isogenic gene-corrected line derived from PD1 (PD1 GC). Replication-induced senescence was achieved through passaging (>15 passages; aged, A), whereas early-passage cells (<8 passages) were defined as young (Y) (Fig. 1a).

Aged NESCs retained the typical three-dimensional colony morphology observed in young cultures (Supplementary Fig. 1a). Immunostaining confirmed the presence of the canonical neural stemness and ventral identity markers SOX2 and FOXA2, respectively, in both young and aged NESCs (Fig. 1b). We performed semi-quantitative analysis of SOX2 and FOXA2 immunostaining. The presence of SOX2 increased in HT and PD1 GC aged NESCs but was unchanged for PD1 and PD2 (Fig. 1d). FOXA2 presence increased in aged HT, PD1, and PD2 NESCs, whereas the PD1 GC line showed the opposite trend (Fig. 1e). To assess whether lineage identity was preserved at the molecular level, we first performed bulk mRNA sequencing. Transcriptomic profiling confirmed expression of stemness (ST), ventral midbrain (VM) patterning, and floor plate (FP) progenitor markers in both young and aged conditions (Fig. 1c), indicating that passaging did not perturb overall lineage commitment. However, when performing differential gene expression, pooling all lines revealed significant downregulation of VM-associated genes in aged NESCs, including *EN1*, *EN2*, *FOXA2* and *OTX2*, alongside alterations in *SOX1* and several HOX genes (Supplementary Table 2), consistent with a shift in regional identity signatures. To investigate these changes more quantitatively, we next scored each sample against a literature-curated gene marker set comprising markers of mitochondria, lipid, and senescence. This analysis revealed line-dependent shifts in aged NESCs: HT, PD1, and PD2 lines showed increased Stem identity scores and decreased Ventral midbrain identity scores with ageing, whereas the PD1 GC line remained unchanged (Extended Fig. 1b).

Beyond identity-related pathways, we performed functional enrichment analysis to assess which molecular processes were altered in the aged NESCs. Gene ontology (GO) analysis of differentially expressed genes between young and aged NESC identified significant enrichment of pathways related to telomerase localisation and negative regulation of the cell cycle (Fig. 1f), indicating activation of molecular programs consistent with replicative exhaustion. In line with this, aged NESCs exhibited reduced *TERT* expression (Fig. 1g, left graph) and decreased telomerase activity scores calculated using the EXTEND algorithm ^33^ (Fig. 1g, right graph), supporting a role for telomere-related mechanisms in passage-induced replicative senescence in our experimental model. In agreement with enrichment results indicating suppression of cell cycle-related pathways, quantified immunostaining for Ki-67 demonstrated a significant reduction in the percentage of proliferative cells across all aged conditions compared to their young counterparts (Fig. 1h, i). We also observed a decreased percentage of cleaved caspase-3-positive cells in aged NESC compared to the young, except for PD2 (Fig. 1h, j), suggesting a potential increased resistance to apoptosis as observed in senescent-like states ^12^.

We next evaluated the gene expression of *GLB1*, encoding for β-galactosidase (Fig. 1k), and *LMNB1* (Fig. 1l), classical markers associated with senescence. Aged NESC showed significantly increased expression of *GLB1* and reduced expression of *LMNB1* compared to young NESCs, consistent with a profile of senescence-like state. We also evaluated the area of LAMNB1 staining in the NESC and observed a decrease for the aged PD1 GC and PD1, while this parameter was unchanged for HT and PD2, suggesting this feature was not effectively achieved in all the lines due to passaging (Supplementary Fig. 1c-d).

Given the established link between mitochondrial dysfunction, ageing, and PD, we next evaluated mitochondrial-related gene expression programs through gene-set scoring. This analysis revealed line-specific shifts: Mitochondrion organisation and Response to oxidative stress signatures were upregulated in aged HT NESCs but downregulated in PD1, with no significant change in PD1 GC or PD2. Mitophagy scores were increased exclusively in aged HT NESCs, while PD1, PD1 GC, and PD2 showed no significant change (Supplementary Fig. 1b). To assess whether these transcriptional changes were accompanied by morphological alterations, we performed immunostaining for TOM20, an outer mitochondrial membrane protein (Supplementary Fig. 1e). Aged NESCs across all lines exhibited a reduced TOM20-positive area compared to young NESCs (Extended Fig. 1f). Interestingly, in HT, PD1 GC, and PD1, but not PD2, mitochondria appeared more arborised in aged compared to young NESCs (Supplementary Fig. 1g).

We also evaluated markers that could suggest increased activation of DNA damage response (DDR) pathways and performed immunofluorescence analysis of phosphorylated γH2AX (p-γH2AX) and 53BP1 foci. The number of p-γH2AX foci per nucleus increased after the passaging procedure, in HT and PD2, while PD1 and PD1 GC were unchanged (Fig. 1m, n). On the other hand, the number of 53BP1 foci per nucleus increased in aged NESCs from all the lines (Fig. 1o, p).

Collectively, these data indicate that passage-induced replicative ageing of NESCs recapitulates certain cellular features commonly associated with senescence. Overall, NESC passaging caused increased FOXA2 expression, reduced proliferation, altered mitochondrial morphology in all the lines, activations of certain DDR-associated markers. Other features, such as nuclear laminar disruption, showed more variable penetrance across the lines, suggesting that the acquisition of the full senescent-like stage might be modulated by genetic background.

### Generation of astrocytes with senescence-associated features

After assessing the effect of passage-induced senescence in NESCs, the effect of this approach was investigated on astrocytes. For this purpose, HT and PD2 hiPSC lines were differentiated into NESCs and subsequently patterned into neural stem cells (NSCs) (Fig. 2a). Replicative aging was induced at this stage through extended passaging and the NSCs were then differentiated into astrocytes (Fig. 2a). Expression of the canonical markers GFAP and S100B confirmed the preservation of the astrocytic identity in aged astrocytes (Fig. 2b). Semi-quantitative image analysis revealed line-specific morphological remodeling: nuclear area decreased in PD2 aged versus the young counterpart while this was unchanged when comparing HT young and aged (Supplementary Fig. 2a). GFAP signal decreased in aged HT but increased in PD2, while arborization (expressed as branching) showed the opposite pattern, increased in aged HT and reduced in aged PD2 (Supplementary Fig. 2b, c).

We performed immunohistochemical staining for cleaved caspase-3-positive area per nucleus area, which was significantly reduced in aged astrocytes compared to their young counterparts in both HT and PD2 (Fig. 2c).

We next performed senescence-associated β-galactosidase (SA-β-gal) activity staining, which showed a significantly increased staining in both HT and PD2 aged lines compared to the young counterparts (Fig. 2d, e). In contrast to NESCs, the Lamin B1 staining showed a significant decrease in expression of the protein in aged astrocytes compared to young astrocytes, consistent with nuclear lamina thinning observed during senescence.

To determine whether these changes were associated with activation of senescence-related stress pathways, DDR signalling was again assessed. Compared to young astrocytes, aged astrocytes displayed significantly increased accumulation of γH2AX (Fig. 2h, i), accompanied by a significantly increased 53BP1 signal (Fig. 2j, k), indicating sustained DDR activation. Expression of the cyclin-dependent kinase inhibitor p21, a downstream effector of DDR-mediated cell cycle arrest, was significantly elevated only in aged PD2 astrocytes compared to young astrocytes (Fig. 2l, m), while its expression was unchanged in HT.

Since both senescence and PD are closely associated with mitochondrial metabolic and morphological changes, we performed a further characterisation. Immunostaining for the outer mitochondrial membrane protein TOM20 revealed an overall significant increase in mitochondrial area in aged compared to young astrocytes (Fig. 2n, o), indicative of increased mitochondrial mass. Mitochondrial morphological analysis revealed increased mitochondrial branching in aged astrocytes compared to young astrocytes (Fig. 2p). Despite enhanced branching, mitochondrial structures appeared more round in aged versus young astrocytes, as evidenced by a significantly increased shape to surface index (Fig. 2q), consistent with more circular and fragmented mitochondria also associated with ageing neurons ^35,36^. To determine whether these structural alterations were accompanied by changes in mitochondrial turnover, mitophagy was assessed using astrocytes expressing the ATP5F1C-Rosella reporter (Fig. 2r). Aged HT astrocytes displayed the same mitophagy rate (Fig. 2s) compared to young HT astrocytes. In contrast, aged PD2 astrocytes exhibited significantly reduced mitophagy (Fig. 2s), in line with previous observation connecting impaired mitophagy with several PD-associated mutations, including LRRK2-G2019S ^37^. These observations suggest changes in mitochondrial morphology due to passaging but decreased mitophagy only in cells with the PD background, suggesting susceptibility due to the LRRK2-G2019S mutation.

Collectively, these data demonstrate that passage-induced replicative senescence at the NSCs stage allows the generation of astrocytes exhibiting features associated with cellular senescence, in particular, increased γH2AX and 53BP1, and mitochondrial roundness.

### Passage-induced ageing at the NESC stage allows for the generation of hMOs with senescence-associated features

To investigate the effects of passage-induced ageing within a multicellular midbrain context, we generated hMOs from young and aged NESCs derived from HT, PD1 GC, PD1, and PD2 iPSC lines (Fig. 3a). hMOs were collected after 70 days of differentiation, a stage at which major midbrain cell populations, including astrocytes and dopaminergic neurons, are present ^30,38^.

Aged hMOs displayed GFAP-positive astrocytes and TH-positive dopaminergic neurons (Fig. 3b). Of note, nuclear area decreased in aged HT and PD2 but was unchanged in PD1 GC and PD1 compared to their young counterparts (Fig. 3c). In parallel, PD1 GC, PD2, and with a similar trend HT aged hMOs showed an increased pyknotic nuclear area, whereas PD1 did not (Supplementary Fig. 3a).

Semi-quantitative analysis of immunostaining revealed line-specific changes in cellular composition. TH positive area was unchanged across all lines except PD2, where it was reduced (Fig. 3d). TH arborization was increased in aged PD2 hMOs, with a similar but non-significant trend in HT, while PD1 GC and PD1 were unchanged (Supplementary Fig. 3b). TH fragmentation was increased in PD2, reduced in HT and PD1 GC, and unchanged in PD1 (Supplementary Fig. 3c). Interestingly, *TH* mRNA levels were significantly increased in HT and PD2, with a similar trend in PD1 but not in PD1 GC (Supplementary Fig. 1d).

GFAP-positive area was significantly decreased in aged PD1 and PD2, significantly increased in PD1 GC, while HT remained unchanged (Fig. 3e). Despite the reduced astrocyte presence in PD1 and PD2, the remaining astrocytes appeared significantly more branched in HT, PD1 and PD2, with no significant change in PD1 GC (Fig. 3f), potentially suggesting a shift toward an activated phenotype.

Following, senescence-associated features connected with nuclear lamina reorganisation and DDR were evaluated in hMOs. *LMNB1* mRNA levels were significantly reduced in aged HT and PD2 hMOs, with a similar but nonsignificant decrease in PD1 GC and PD1 (Fig. 3g). DDR activation was assessed specifically in astrocytes within hMOs. Immunohistochemical analysis revealed significantly increased expression of γH2AX in astrocytic nuclei in aged HT and PD2 hMOs, with a similar but nonsignificant trend in PD1 and PD1 GC (Fig. 3 h, i). A similar trend was observed for 53BP1 foci formation: increased in aged HT, PD1 GC, and PD2 hMOs, and non-significant increases for PD1 hMOs (Fig. 3j, k), paralleling the γH2AX pattern and further supporting enhanced DDR signalling. At the transcript level, HMGB1, a chromatin-associated protein whose loss from the nucleus is a hallmark of senescence, showed unaltered mRNA levels across all lines (Supplementary Fig. 3e). Similarly, TP53BP1 mRNA was upregulated only in PD2, with no significant changes in HT, PD1 GC, or PD1 (Supplementary Fig. 3f).

Together, these data indicate that passage-induced senescence extends to the hMOs context, although genotype-specific differences are apparent. Notably, the data also reveal that the genetic background of each cell line might actively modulate the manifestation of these hallmarks under the passaging procedure.

### Passage-induced alteration of lipid homeostasis in hMOs

A central feature of brain ageing is substantial alterations in lipid composition ^39^. To assess this in the hMOs, we performed global lipidome profiling. The analysis revealed widespread remodelling, assessed by expression profiles between young and aged hMOs of the same line (Fig. 4a; Supplementary Fig. 4a-c). Differential abundance analysis demonstrated an overall reduction in several lipid classes across three of four lines, except for PD1 GC, including lysophosphatidylcholines (LPC), plasmalogen LPC (P-LPC), sphingomyelins (SM), glucosylceramides (GlcCer), and acylcarnitines (CAR). Cholesteryl esters (CE) represented the only lipid class consistently reduced across all cell lines. Lipid ontology (LION) analysis revealed that, in aged HT, PD1 and PD2 hMOs, endoplasmic reticulum-associated lipid species were enriched (Fig. 4b), consistent with the central role of the ER in lipid biosynthesis and membrane remodelling. In aged PD1 and PD2 hMOs, we observed additional enrichment for mitochondrial-associated lipid species (Fig. 4b, bottom panel), consistent with the mitochondrial alterations seen in ageing and LRRK2 PD ^40^.

To further characterise coordinated lipid alterations, weighted lipid co-expression network analysis (WGCNA) was applied to identify lipid co-abundance modules (Supplementary Fig. 4d). Module eigengenes (MEs) were modelled using linear regression (ME ∼ Line × Ageing) (Fig. 4c; Supplementary Fig. 4e). The generated ageing coefficient (β Ageing) represents the effect of ageing across genotypes after adjustment for baseline differences between lines. Modules ME1 (turquoise), ME2 (blue), and ME4 (yellow) exhibited significant positive ageing coefficients (β > 0, FDR < 0.01), indicating increased module-level lipid abundance in aged hMOs. In contrast, ME3 (brown) showed a significant negative ageing coefficient (β < 0, FDR < 0.01), consistent with reduced lipid abundance in aged conditions.

A significant Line × Ageing interaction term for ME1, ME3, and ME4 (FDR < 0.05) indicates that although the direction of the ageing effect was generally consistent, its magnitude varied across genotypes. This genotype-dependent modulation became evident when examining module-level lipid abundance stratified by line. For example, ME3 displayed a higher baseline abundance in PD1 and PD2 compared to HT, whereas the PD1 GC baseline is comparable to HT (Supplementary Fig. 4e). These data indicate that differences between cell lines modulate the magnitude of the changes, while the direction of the effects remains consistent.

Module composition analysis revealed that more than 50% of lipid species within the strongest ageing-associated modules ME1 and ME3 belonged to glycerophosphocholines (PC), glycerophosphoethanolamines (PE), and triacylglycerols (TG) (Fig. 4d, Extended Fig. 3f). To identify central lipid drivers within each module, module eigengene-based connectivity (kME) was calculated. Lipids with high absolute kME values represent species whose abundance closely tracks the dominant module signal. Notably, several PC and triacylglycerols (TG) species were prominent drivers within ME1, whereas ME3 was enriched for PC and PE. Disruption of Glycerophospholipids, in particular LPC and TG, was reported in the PD brain ^41^.

Together, these findings showed that passage-induced ageing in hMOs recapitulates coordinated remodelling of lipid species abundance, encompassing both global and genotype-dependent effects.

### PD reshapes passage-induced senescence programs in a category- and cell-type-dependent manner

To obtain a systems-level view of the biological programs altered by passage-induced senescence across cell types and genetic backgrounds, we computed category-level scores as the mean z-score of all features assigned to each biological Category × Modality combination using a linear model fitted per Category, and examined across NESCs, astrocytes, and hMOs (Fig. 5a; Supplementary Fig. 5a, b, c for the individual features). In NESCs, several Senescence Hallmarks categories, including Apoptosis, Cell Cycle, and DDR, were altered regardless of genotype. Mitochondrial Homeostasis categories, including Mitophagy and Autophagy gene set, increased with passaging in HT and GC NESCs but were attenuated or had an opposite trend in PD lines. In astrocytes, a similar pattern was observed: Senescence, DDR, and Mitochondrial Content, while Apoptosis and Nuclear Lamina decreased. Mitophagy was elevated with passaging in HT astrocytes but showed opposite trajectories in PD lines. In hMOs, Lipid Remodelling was reduced across HT and PD genotypes but not in GC one, except for Sterol Lipids, which are consistently reduced, and Nuclear Lamina scores are also decreased irrespective of genetic background. DaN Fragmentation diverged between genotypes, increasing with passaging in PD hMOs while decreasing in HT and GC, potentially suggesting an early signature of genotype-dependent neuronal vulnerability.

To determine whether the PD genetic background imposes an altered baseline state, PD young category scores were compared to HT young and HT aged references across NESCs, astrocytes and hMOs (Fig. 5b, c). In NESCs, PD young cells exhibited elevated scores for Senescence, Mitochondrial Function, Lipid Remodelling, and Mitophagy relative to the HT young baseline, along with reduced scores for Telomere, Chromatin State, Ventral Midbrain Identity and Apoptosis (Supplementary Fig. 5b). These patterns indicate that PD young cells already resemble senescence-associated changes observed in HT aged NESCs relative to the HT young baseline for several Categories. In astrocytes, PD young cells also showed DDR, Senescence, Astrocyte Ramification, and Mitochondrial Content Category scores comparable to those of HT aged cells in respect of HT young baseline (Fig. 5b). Conversely, Astrocytes Features and Apoptosis categories, which increased in aged HT astrocytes, were reduced in PD young cells, demonstrating that this premature senescent phenotype is selective for certain features rather than a uniform shift toward an aged state. In hMOs, PD young hMOs displayed DDR category scores comparable to HT aged hMOs (Fig. 5c). All lipid classes except for Glycerophospholipids, as well as Nuclear Lamina and Nuclear Area categories that decline with passaging in HT hMOs, were likewise reduced in PD young hMOs, consistent with a convergent trajectory in which the PD genetic background recapitulates aged lipid and nuclear architecture alterations similarly to HT aged, and prematurely relative to HT young baseline.

Beyond baseline differences, we assessed whether the PD genetic background modulates senescent-like phenotypes and assessed whether senescence-associated Category changes differ in magnitude or direction between PD and HT lines. In NESCs, Mitophagy, Mitochondrial Function, Lipid Remodeling, and Nuclear Lamina (ICC) Categories showed decreased magnitude of senescence-associated changes in PD lines compared to HT, while Chromatin State and, paradoxically, Nuclear Lamina (RNA) enhanced senescence-associated increase in PD NESCs (Supplementary Fig. 5c). In astrocytes, Astrocyte Features and Nuclear Lamina showed exacerbated aging-associated changes in PD relative to HT (Fig. 5d). In contrast, Mitophagy exhibited attenuated aging-associated changes in PD astrocytes, although not significant, indicating that the PD genetic background selectively constrains mitochondrial quality control responses during aging. In hMOs, DaN Fragmentation change is significantly exacerbated during passaging in PD than in HT hMOs, while DDR (qPCR) Category displayed reduced age-related changes in PD (Fig. 5e).

Together, these findings suggest that the PD genetic background accelerates the rate of acquisition of certain senescent-like features in a category- and cell-type-dependent manner. Notably, senescence in PD hMOs amplified DaN Fragmentation and Apoptosis, key pathological features of PD, suggesting that the combined effect of the PD genetic background and senescence exacerbates PD-relevant neuronal vulnerability in the context of LRRK2-G2019S in our system.

## Discussion

Extended passaging of NSCs and NESCs induced molecular and cellular alterations that recapitulated certain features of cellular senescence, including reduced proliferative capacity, nuclear lamina disruption, and DNA damage response (DDR) activation. These features were maintained upon differentiation into astrocytes and were partially retained within hMOs, where they co-occurred with lipidomic remodelling and a selective reduction in dopaminergic neuron content in PD genetic backgrounds. We observed that the LRRK2-G2019S background modulated a few senescence-associated features under replicative exhaustion conditions, such as reduction of mitophagy in astrocytes or enrichment of mitochondrial-associated lipid species in hMOs. Senescence and degeneration are clearly extremely complex and intersecting processes. However, our data suggest that some aspects of this complex process can be recapitulated *in vitro* by using patient-specific cells and inducing replicative senescence.

Replicative exhaustion represents an established approach to induce senescence-associated states *in vitro*^42^. Strategies based on acute stressors, irradiation, or progeroid gene expression are effective tools. However, they might add confounders, which are difficult to account for when studying diseases such as PD. Extended passaging engages endogenous cell-intrinsic programs associated with replicative decline. Replicative senescence is mechanistically distinct from acute stress-induced senescence and produces partially overlapping but non-identical molecular phenotypes^43^. Astrocytes are known to undergo replicative senescence in culture^44–46^. The present findings indicate that similar processes can be harnessed in human iPSC-derived midbrain progenitors (*e.g.* NSCs and NESCs) while preserving differentiation competence. Importantly, phenotypes were observed in differentiated astrocytes and hMOs, enabling investigation of senescence-associated programs within a multicellular human system.

Among the most prominent features observed in astrocytes derived from high-passage progenitors was mitochondrial remodelling. Mitochondrial alterations are established hallmarks of both cellular senescence and physiological ageing. Senescent cells frequently compensate for mitochondrial dysfunction through increased mitochondrial biogenesis, leading to elevated mitochondrial mass that is often accompanied by reduced mitochondrial turnover or impaired mitophagy. We found altered mitochondrial morphology, increased organelle mass, and genotype-dependent changes in mitophagy were evident following replicative exhaustion in aged NESCs (Supplementary Fig.1 f, g) and aged astrocytes (Fig. 2n -s, Supplementary Fig. 2 d). Aged astrocytes exhibited increased mitochondrial mass (Fig. 2o) and a more ramified network (Fig. 2p), yet individual mitochondrial bodies appeared more rounded (Fig. 2q), as indicated by an increased shape-to-surface index. This combination, network-level branching alongside individual mitochondria circularity, is consistent with the mitochondrial remodelling reported in senescent astrocytes, where fragmentation of individual mitochondria occurs against a background of elevated biogenesis and impaired fission-fusion balance^47^. In PD, these structural alterations were accompanied by reduced mitophagy rates (Fig. 2s, PD2). Mitochondrial dysfunction and mitophagy defects are likewise central features of PD pathogenesis, particularly in the context of the LRRK2-G2019S mutation^28,48^. Impaired mitophagy and increased reactive oxygen species can reinforce persistent DDR signalling, which can engage mitochondrial pathways that stabilise and sustain senescence-associated programs^49,50^. Within this framework, aged HT astrocytes show an increase in mitochondrial mass accompanied by a slight increase in mitophagy (Fig. 2s, HT). In contrast, LRRK2-G2019S astrocytes exhibit reduced mitophagy despite increased mitochondrial mass reduced mitophagy (Fig. 2s, PD2), suggesting that the PD-associated genetic mutation amplifies senescence-related dysregulation of mitochondrial homeostasis. Given the central role of mitochondrial homeostasis in PD, this genotype-dependent enhancement of the senescence-mitochondria axis may represent a mechanistic point of convergence between aging-related cellular programs and inherited disease susceptibility, as also revealed by the category level analysis (Fig. 5d). Interestingly, the mitophagy category in PD young samples is more similar to the HT aged condition than to HT young, indicating that this process is already altered in PD per se and is further exacerbated in aged PD astrocytes (Fig. 5b).

Beyond mitochondrial alterations, hMOs derived from high-passage progenitors exhibited lipidomic remodelling compared to those generated from low-passage progenitors (Fig. 4) in both HT and PD cell lines, although this was not observed in the PD1 GC line. Brain ageing is known to be accompanied by coordinated shifts in lipid composition^39,51,52^, and lipid dysregulation is increasingly recognised as a feature of neurodegenerative disorders, including Alzheimer’s disease^53^ and PD^41^. The convergence of DDR activation, mitochondrial remodelling, and lipid alterations in our model suggests that these processes are interconnected. Mitochondrial dysfunction in senescent glia has been linked to altered lipid handling and accumulation in Drosophila ^54^. Mitochondrial impairment in PD is likewise associated with lipid dysregulation ^40^, supporting the idea that coordinated disruption of these pathways may be pathophysiologically relevant. The occurrence of lipid alterations in hMOs and the observation that in aged astrocytes, mitochondrial homeostasis is altered, therefore might indicate broader metabolic reprogramming accompanying astrocyte senescence.

A recurring observation across our data is that individual senescence-associated markers show variable penetrance across cell lines and, in some cases, across cell types. For example, inflammatory readouts such as *IL6* and *IL1B* exhibited heterogeneous responses across PD lines, and certain markers did not reach consistent significance in hMOs contexts (*e.g. HMGB1*, *p16/CDKN2A*). This phenomenon is well-documented in the senescence literature^55–57^. Individual canonical readouts such as p16/CDKN2A, HMGB1, and inflammatory SASP components are not universally or consistently induced across cell types, genetic backgrounds, or senescence-inducing stimuli^43^. This heterogeneity, which has also been called "senotype", reflects the distinct combinations of molecular programs adopted by senescent cells depending on their context and the potential dynamic range of this process^43^. This means that single-marker or single-modality assessments are likely limited in capturing whether a given cell population has entered a senescence-like state^58^. Considering this aspect, a key observation of our multi-modal integrative analysis is that, despite the line-to-line variability in individual features, the primary discriminant emerging from the category-level heatmaps (Fig. 5a, Supplementary Fig. 5a-c) is passage condition: young versus aged, rather than genetic background. Samples within the same passage condition cluster together across cell lines, indicating that, while individual senescence hallmarks are variably penetrant, the collective, multi-modal signature of replicative exhaustion is sufficiently coherent and reproducible to overcome inter-line differences at the level of biological categories. Our findings, therefore, support the view that multi-modal integration, also in iPSC-derived models, combining morphometric, transcriptomic, and lipidomic readouts, is necessary to capture complex and potentially subtle, senescent-associated changes.

Further studies will be essential to validate the findings in this study, including starting from a higher cohort of iPSC cell lines. The baseline described here relies on a single HT iPSC line. Given the well-documented inter-line variability inherent to iPSC biology, replication with additional healthy donor lines would reinforce the conclusions of the integrative analysis.

Moreover, while passage-induced replicative exhaustion is a physiologically established approach to modelling cellular ageing, it produces a senescence-associated state that might be molecularly distinct from the full replicative senescence achieved in primary somatic cells and from the complex, heterogeneous senescent populations found in the ageing human brain. Not all canonical senescence markers were consistently induced across all lines or cell types: *e.g.* nuclear lamina disruption showed variable penetrance in NESCs. These findings suggest that the model captures a senescence-like state rather than fully recapitulating the complete senescence program, and that certain hallmarks may require additional passages, complementary stimuli, or longer differentiation periods to manifest robustly.

The model presented here offers a human system in which certain senescence-associated cellular programs can be studied in conjunction with genetic susceptibility, in our case, PD. By incorporating replicative exhaustion upstream of lineage specification, this approach enables the generation of astrocytes, in 2D and hMOs, containing astrocytes presenting senescence-like markers without reliance on acute stressors or progeroid manipulation that could act as confounders to detect truly disease-associated mechanisms. This model can thus be used to investigate the interplay between astrocyte senescence and genotype susceptibility to PD hallmarks. In addition, the platform may facilitate future investigation of interventions targeting senescence-associated pathways with senolytics or senomorphics, for example, in a human midbrain context.

## Research funding

DF was supported by the FNR Core C21/DM/15839823. VC was supported by the TKI MUMC/FHML LSHM202414, 24.0937 project (Health Holland). NYL was supported by the Starting Grant FHML 2024, 25.1117.

## Contributions

SB conceived and designed the study; SB obtained funding. VC, NLY and DF performed the cell culture experiments; VC and NLY performed the NESCs immunostaining (Fig.1, Supplementary Fig.1, Fig 5, Supplementary Fig.5); DF performed the astrocytes immunostaining and live cell imaging (Fig.2, Supplementary Fig.2, Fig.5), VC performed the hMOs immunostaining (Fig.3, Fig.5, Supplementary Fig.5); DF performed the SA-β-gal activity assay (Fig.2, Fig.5, Supplementary Fig.5); VC performed the hMOs qPCR (Fig.3, Supplementary Fig.3, Fig.5, Supplementary Fig.5); NLY generated the code for the analysis of the NESCs immunostaining (Fig.1, Supplementary Fig.1, Fig 5, Supplementary Fig.5);. NLY analyzed the NESCs immunostaining (Fig.1, Supplementary Fig.1, Fig 5, Supplementary Fig.5); DF analyzed the astrocytes immunostaining, live cell, and SA-β-gal activity assay (Fig2., Supplementary Fig.2); AZ analyzed the hMOs immunostaining (Fig.3, Supplementary Fig.3, Fig.5, Supplementary Fig.5); VC analyzed the RNA-sequencing data of the NESCs (Fig.1, Supplementary Fig.1, Fig5, Supplementary Fig.5); VC analyzed the lipidomic data of the hMOs (Fig.4, Supplementary Fig.4, Fig5, Supplementary Fig.5); SB and VC designed the multi-modal analysis (Fig.5, Supplementary Fig.5). JCS and JJ provided the mitophagy lines, analysed and interpreted the data (Fig.2 q-s; Supplementary Fig.2). SB and VC interpreted the data. VC generated the code for all the graphs and prepared all the figures. VC drafted the manuscript with input and technical description from all the authors; all authors reviewed and approved the manuscript.

## References

1. Collier, T.J., Kanaan, N.M., and Kordower, J.H. (2011). Ageing as a primary risk factor for Parkinson’s disease: evidence from studies of non-human primates. Nat Rev Neurosci 12, 359–366. 10.1038/nrn3039.

2. Schmidt, M.Y., Cuervo, A.M., Double, K.L., Ehninger, D., Goldberg, M.S., Harvey, K., Hoeijmakers, J.H.J., Luk, K.C., Mastroberardino, P.G., Moore, D.J., et al. (2026). Unraveling the intersection of aging and Parkinson’s disease: a collaborative roadmap for advancing research models. NPJ Parkinsons Dis 12, 28. 10.1038/s41531-025-01239-x.

3. Morris, H.R., Spillantini, M.G., Sue, C.M., and Williams-Gray, C.H. (2024). The pathogenesis of Parkinson’s disease. Lancet 403, 293–304. 10.1016/S0140-6736(23)01478-2.

4. Zimprich, A., Biskup, S., Leitner, P., Lichtner, P., Farrer, M., Lincoln, S., Kachergus, J., Hulihan, M., Uitti, R.J., Calne, D.B., et al. (2004). Mutations in LRRK2 cause autosomal-dominant parkinsonism with pleomorphic pathology. Neuron 44, 601–607. 10.1016/j.neuron.2004.11.005.

5. Surmeier, D.J. (2018). Determinants of dopaminergic neuron loss in Parkinson’s disease. FEBS J 285, 3657–3668. 10.1111/febs.14607.

6. Kim, H.Y., Kim, S., Akaydin, A.N., Kim, S., Hyeon, S.J., Lee, J., and Ryu, H. (2026). The rise of astrocytes: are they guardians or troublemakers of the brain disorder? Exp Mol Med 58, 301–318. 10.1038/s12276-025-01627-6.

7. Lee, H.G., Wheeler, M.A., and Quintana, F.J. (2022). Function and therapeutic value of astrocytes in neurological diseases. Nat Rev Drug Discov 21, 339–358. 10.1038/s41573-022-00390-x.

8. Ramos-Gonzalez, P., Mato, S., Chara, J.C., Verkhratsky, A., Matute, C., and Cavaliere, F. (2021). Astrocytic atrophy as a pathological feature of Parkinson’s disease with LRRK2 mutation. NPJ Parkinsons Dis 7, 31. 10.1038/s41531-021-00175-w.

9. Sonninen, T.M., Hamalainen, R.H., Koskuvi, M., Oksanen, M., Shakirzyanova, A., Wojciechowski, S., Puttonen, K., Naumenko, N., Goldsteins, G., Laham-Karam, N., et al. (2020). Metabolic alterations in Parkinson’s disease astrocytes. Sci Rep 10, 14474. 10.1038/s41598-020-71329-8.

10. di Domenico, A., Carola, G., Calatayud, C., Pons-Espinal, M., Munoz, J.P., Richaud-Patin, Y., Fernandez-Carasa, I., Gut, M., Faella, A., Parameswaran, J., et al. (2019). Patient-Specific iPSC-Derived Astrocytes Contribute to Non-Cell-Autonomous Neurodegeneration in Parkinson’s Disease. Stem Cell Reports 12, 213–229. 10.1016/j.stemcr.2018.12.011.

11. Muñoz-Espín, D., and Serrano, M. (2014). Cellular senescence: from physiology to pathology. Nature Reviews Molecular Cell Biology 15, 482–496. 10.1038/nrm3823.

12. Hernandez-Segura, A., Nehme, J., and Demaria, M. (2018). Hallmarks of Cellular Senescence. Trends Cell Biol 28, 436–453. 10.1016/j.tcb.2018.02.001.

13. Muwanigwa, M.N., Modamio-Chamarro, J., Antony, P.M.A., Gomez-Giro, G., Kruger, R., Bolognin, S., and Schwamborn, J.C. (2024). Alpha-synuclein pathology is associated with astrocyte senescence in a midbrain organoid model of familial Parkinson’s disease. Mol Cell Neurosci 128, 103919. 10.1016/j.mcn.2024.103919.

14. Chinta, S.J., Woods, G., Demaria, M., Rane, A., Zou, Y., McQuade, A., Rajagopalan, S., Limbad, C., Madden, D.T., Campisi, J., and Andersen, J.K. (2018). Cellular Senescence Is Induced by the Environmental Neurotoxin Paraquat and Contributes to Neuropathology Linked to Parkinson’s Disease. Cell Rep 22, 930–940. 10.1016/j.celrep.2017.12.092.

15. Pertusa, M., Garcia-Matas, S., Rodriguez-Farre, E., Sanfeliu, C., and Cristofol, R. (2007). Astrocytes aged in vitro show a decreased neuroprotective capacity. J Neurochem 101, 794–805. 10.1111/j.1471-4159.2006.04369.x.

16. Kawano, H., Katsurabayashi, S., Kakazu, Y., Yamashita, Y., Kubo, N., Kubo, M., Okuda, H., Takasaki, K., Kubota, K., Mishima, K., et al. (2012). Long-term culture of astrocytes attenuates the readily releasable pool of synaptic vesicles. PLoS One 7, e48034. 10.1371/journal.pone.0048034.

17. Liu, G.H., Barkho, B.Z., Ruiz, S., Diep, D., Qu, J., Yang, S.L., Panopoulos, A.D., Suzuki, K., Kurian, L., Walsh, C., et al. (2011). Recapitulation of premature ageing with iPSCs from Hutchinson-Gilford progeria syndrome. Nature 472, 221–225. 10.1038/nature09879.

18. Vera, E., Bosco, N., and Studer, L. (2016). Generating Late-Onset Human iPSC-Based Disease Models by Inducing Neuronal Age-Related Phenotypes through Telomerase Manipulation. Cell Rep 17, 1184–1192. 10.1016/j.celrep.2016.09.062.

19. Harley, J., Santosa, M.M., Ng, C.Y., Grinchuk, O.V., Hor, J.H., Liang, Y., Lim, V.J., Tee, W.W., Ong, D.S.T., and Ng, S.Y. (2024). Telomere shortening induces aging-associated phenotypes in hiPSC-derived neurons and astrocytes. Biogerontology 25, 341–360. 10.1007/s10522-023-10076-5.

20. Fathi, A., Mathivanan, S., Kong, L., Petersen, A.J., Harder, C.R.K., Block, J., Miller, J.M., Bhattacharyya, A., Wang, D., and Zhang, S.C. (2022). Chemically induced senescence in human stem cell-derived neurons promotes phenotypic presentation of neurodegeneration. Aging Cell 21, e13541. 10.1111/acel.13541.

21. Oyefeso, F.A., Goldberg, G., Opoku, N., Vazquez, M., Bertucci, A., Chen, Z., Wang, C., Muotri, A.R., and Pecaut, M.J. (2023). Effects of acute low-moderate dose ionizing radiation to human brain organoids. PLoS One 18, e0282958. 10.1371/journal.pone.0282958.

22. Shakhbazau, A., Danilkovich, N., Seviaryn, I., Ermilova, T., and Kosmacheva, S. (2019). Effects of minocycline and rapamycin in gamma-irradiated human embryonic stem cells-derived cerebral organoids. Mol Biol Rep 46, 1343–1348. 10.1007/s11033-018-4552-6.

23. Arias-Fuenzalida, J., Jarazo, J., Walter, J., Gomez-Giro, G., Forster, J.I., Krueger, R., Antony, P.M.A., and Schwamborn, J.C. (2019). Automated high-throughput high-content autophagy and mitophagy analysis platform. Scientific Reports 9. 10.1038/s41598-019-45917-2.

24. Burridge, P.W., Thompson, S., Millrod, M.A., Weinberg, S., Yuan, X., Peters, A., Mahairaki, V., Koliatsos, V.E., Tung, L., and Zambidis, E.T. (2011). A Universal System for Highly Efficient Cardiac Differentiation of Human Induced Pluripotent Stem Cells That Eliminates Interline Variability. PLoS ONE 6, e18293. 10.1371/journal.pone.0018293.

25. Reinhardt, P., Glatza, M., Hemmer, K., Tsytsyura, Y., Thiel, C.S., Hoing, S., Moritz, S., Parga, J.A., Wagner, L., Bruder, J.M., et al. (2013). Derivation and expansion using only small molecules of human neural progenitors for neurodegenerative disease modeling. PLoS One 8, e59252. 10.1371/journal.pone.0059252.

26. Reinhardt, P., Schmid, B., Burbulla, L.F., Schondorf, D.C., Wagner, L., Glatza, M., Hoing, S., Hargus, G., Heck, S.A., Dhingra, A., et al. (2013). Genetic correction of a LRRK2 mutation in human iPSCs links parkinsonian neurodegeneration to ERK-dependent changes in gene expression. Cell Stem Cell 12, 354–367. 10.1016/j.stem.2013.01.008.

27. Nickels, S.L., Walter, J., Bolognin, S., Gerard, D., Jaeger, C., Qing, X., Tisserand, J., Jarazo, J., Hemmer, K., Harms, A., et al. (2019). Impaired serine metabolism complements LRRK2-G2019S pathogenicity in PD patients. Parkinsonism Relat Disord 67, 48–55. 10.1016/j.parkreldis.2019.09.018.

28. Walter, J., Bolognin, S., Antony, P.M.A., Nickels, S.L., Poovathingal, S.K., Salamanca, L., Magni, S., Perfeito, R., Hoel, F., Qing, X., et al. (2019). Neural Stem Cells of Parkinson’s Disease Patients Exhibit Aberrant Mitochondrial Morphology and Functionality. Stem Cell Reports 12, 878–889. 10.1016/j.stemcr.2019.03.004.

29. Palm, T., Bolognin, S., Meiser, J., Nickels, S., Trager, C., Meilenbrock, R.L., Brockhaus, J., Schreitmuller, M., Missler, M., and Schwamborn, J.C. (2015). Rapid and robust generation of long-term self-renewing human neural stem cells with the ability to generate mature astroglia. Sci Rep 5, 16321. 10.1038/srep16321.

30. Monzel, A.S., Smits, L.M., Hemmer, K., Hachi, S., Moreno, E.L., van Wuellen, T., Jarazo, J., Walter, J., Bruggemann, I., Boussaad, I., et al. (2017). Derivation of Human Midbrain-Specific Organoids from Neuroepithelial Stem Cells. Stem Cell Reports 8, 1144–1154. 10.1016/j.stemcr.2017.03.010.

31. Matyash, V., Liebisch, G., Kurzchalia, T.V., Shevchenko, A., and Schwudke, D. (2008). Lipid extraction by methyl-tert-butyl ether for high-throughput lipidomics. J Lipid Res 49, 1137–1146. 10.1194/jlr.D700041-JLR200.

32. Breitkopf, S.B., Ricoult, S.J.H., Yuan, M., Xu, Y., Peake, D.A., Manning, B.D., and Asara, J.M. (2017). A relative quantitative positive/negative ion switching method for untargeted lipidomics via high resolution LC-MS/MS from any biological source. Metabolomics 13. 10.1007/s11306-016-1157-8.

33. Noureen, N., Wu, S., Lv, Y., Yang, J., Alfred Yung, W.K., Gelfond, J., Wang, X., Koul, D., Ludlow, A., and Zheng, S. (2021). Integrated analysis of telomerase enzymatic activity unravels an association with cancer stemness and proliferation. Nat Commun 12, 139. 10.1038/s41467-020-20474-9.

34. Hanzelmann, S., Castelo, R., and Guinney, J. (2013). GSVA: gene set variation analysis for microarray and RNA-seq data. BMC Bioinformatics 14, 7. 10.1186/1471-2105-14-7.

35. Varghese, N., Szabo, L., Cader, M.Z., Lejri, I., Grimm, A., and Eckert, A. (2025). Tracing mitochondrial marks of neuronal aging in iPSCs-derived neurons and directly converted neurons. Commun Biol 8, 723. 10.1038/s42003-025-08152-2.

36. Jiang, H.C., Hsu, J.M., Yen, C.P., Chao, C.C., Chen, R.H., and Pan, C.L. (2015). Neural activity and CaMKII protect mitochondria from fragmentation in aging Caenorhabditis elegans neurons. Proc Natl Acad Sci U S A 112, 8768–8773. 10.1073/pnas.1501831112.

37. Wauters, F., Cornelissen, T., Imberechts, D., Martin, S., Koentjoro, B., Sue, C., Vangheluwe, P., and Vandenberghe, W. (2020). LRRK2 mutations impair depolarization-induced mitophagy through inhibition of mitochondrial accumulation of RAB10. Autophagy 16, 203–222. 10.1080/15548627.2019.1603548.

38. Smits, L.M., Magni, S., Kinugawa, K., Grzyb, K., Luginbuhl, J., Sabate-Soler, S., Bolognin, S., Shin, J.W., Mori, E., Skupin, A., and Schwamborn, J.C. (2020). Single-cell transcriptomics reveals multiple neuronal cell types in human midbrain-specific organoids. Cell Tissue Res 382, 463–476. 10.1007/s00441-020-03249-y.

39. Yu, Q., He, Z., Zubkov, D., Huang, S., Kurochkin, I., Yang, X., Halene, T., Willmitzer, L., Giavalisco, P., Akbarian, S., and Khaitovich, P. (2020). Lipidome alterations in human prefrontal cortex during development, aging, and cognitive disorders. Mol Psychiatry 25, 2952–2969. 10.1038/s41380-018-0200-8.

40. Hallqvist, J., Toomey, C.E., Pinto, R., Baldwin, T., Doykov, I., Wernick, A., Al Shahrani, M., Evans, J.R., Lachica, J., Pope, S., et al. (2025). Multi-omic analysis reveals lipid dysregulation associated with mitochondrial dysfunction in parkinson’s disease brain. Nat Commun 16, 10490. 10.1038/s41467-025-65489-2.

41. Yilmaz, A., Ashrafi, N., Ashrafi, R., Akyol, S., Saiyed, N., Kerseviciute, I., Gabrielaite, M., Gordevicius, J., and Graham, S.F. (2025). Lipid profiling of Parkinson’s disease brain highlights disruption in Lysophosphatidylcholines, and triacylglycerol metabolism. NPJ Parkinsons Dis 11, 159. 10.1038/s41531-025-01023-x.

42. Veronesi, F., Contartese, D., Di Sarno, L., Borsari, V., Fini, M., and Giavaresi, G. (2023). In Vitro Models of Cell Senescence: A Systematic Review on Musculoskeletal Tissues and Cells. Int J Mol Sci 24. 10.3390/ijms242115617.

43. Hernandez-Segura, A., de Jong, T.V., Melov, S., Guryev, V., Campisi, J., and Demaria, M. (2017). Unmasking Transcriptional Heterogeneity in Senescent Cells. Curr Biol 27, 2652–2660 e2654. 10.1016/j.cub.2017.07.033.

44. Kim, D., Yoo, S.H., Yeon, G.B., Oh, S.S., Shin, W.H., Kang, H.C., Lee, C.K., Kim, H.W., and Kim, D.S. (2024). Senescent Astrocytes Derived from Human Pluripotent Stem Cells Reveal Age-Related Changes and Implications for Neurodegeneration. Aging Dis 16, 1709–1731. 10.14336/AD.2024.0089.

45. Blasko, I., Stampfer-Kountchev, M., Robatscher, P., Veerhuis, R., Eikelenboom, P., and Grubeck-Loebenstein, B. (2004). How chronic inflammation can affect the brain and support the development of Alzheimer’s disease in old age: the role of microglia and astrocytes. Aging Cell 3, 169–176. 10.1111/j.1474-9728.2004.00101.x.

46. Evans, R.J., Wyllie, F.S., Wynford-Thomas, D., Kipling, D., and Jones, C.J. (2003). A P53-dependent, telomere-independent proliferative life span barrier in human astrocytes consistent with the molecular genetics of glioma development. Cancer Res 63, 4854–4861.

47. Diniz, L.P., Araujo, A.P.B., Carvalho, C.F., Matias, I., de Sa Hayashide, L., Marques, M., Pessoa, B., Andrade, C.B.V., Vargas, G., Queiroz, D.D., et al. (2024). Accumulation of damaged mitochondria in aging astrocytes due to mitophagy dysfunction: Implications for susceptibility to mitochondrial stress. Biochim Biophys Acta Mol Basis Dis 1870, 167470. 10.1016/j.bbadis.2024.167470.

48. Bolognin, S., Fossépré, M., Qing, X., Jarazo, J., Ščančar, J., Moreno, E.L., Nickels, S.L., Wasner, K., Ouzren, N., Walter, J., et al. (2019). 3D Cultures of Parkinson’s Disease-Specific Dopaminergic Neurons for High Content Phenotyping and Drug Testing. Advanced Science 6, 1800927. 10.1002/advs.201800927.

49. Correia-Melo, C., Marques, F.D., Anderson, R., Hewitt, G., Hewitt, R., Cole, J., Carroll, B.M., Miwa, S., Birch, J., Merz, A., et al. (2016). Mitochondria are required for pro-ageing features of the senescent phenotype. EMBO J 35, 724–742. 10.15252/embj.201592862.

50. Yamauchi, S., Sugiura, Y., Yamaguchi, J., Zhou, X., Takenaka, S., Odawara, T., Fukaya, S., Fujisawa, T., Naguro, I., Uchiyama, Y., et al. (2024). Mitochondrial fatty acid oxidation drives senescence. Sci Adv 10, eado5887. 10.1126/sciadv.ado5887.

51. Tu, J., Yin, Y., Xu, M., Wang, R., and Zhu, Z.J. (2017). Absolute quantitative lipidomics reveals lipidome-wide alterations in aging brain. Metabolomics 14, 5. 10.1007/s11306-017-1304-x.

52. Mota-Martorell, N., Andres-Benito, P., Martin-Gari, M., Galo-Licona, J.D., Sol, J., Fernandez-Bernal, A., Portero-Otin, M., Ferrer, I., Jove, M., and Pamplona, R. (2022). Selective brain regional changes in lipid profile with human aging. Geroscience 44, 763–783. 10.1007/s11357-022-00527-1.

53. Feringa, F.M., Koppes-den Hertog, S.J., Wang, L.Y., Derks, R.J.E., Kruijff, I., Erlebach, L., Heijneman, J., Miramontes, R., Pompner, N., Blomberg, N., et al. (2025). The Neurolipid Atlas: a lipidomics resource for neurodegenerative diseases. Nat Metab 7, 2142–2164. 10.1038/s42255-025-01365-z.

54. Byrns, C.N., Perlegos, A.E., Miller, K.N., Jin, Z., Carranza, F.R., Manchandra, P., Beveridge, C.H., Randolph, C.E., Chaluvadi, V.S., Zhang, S.L., et al. (2024). Senescent glia link mitochondrial dysfunction and lipid accumulation. Nature 630, 475–483. 10.1038/s41586-024-07516-8.

55. Walker, K.A., Basisty, N., Wilson, D.M., 3rd, and Ferrucci, L. (2022). Connecting aging biology and inflammation in the omics era. J Clin Invest 132. 10.1172/JCI158448.

56. Basisty, N., Kale, A., Jeon, O.H., Kuehnemann, C., Payne, T., Rao, C., Holtz, A., Shah, S., Sharma, V., Ferrucci, L., et al. (2020). A proteomic atlas of senescence-associated secretomes for aging biomarker development. PLoS Biol 18, e3000599. 10.1371/journal.pbio.3000599.

57. Anerillas, C., Altes, G., Gresova, K., Tsitsipatis, D., Mazan-Mamczarz, K., Banarjee, R., Cunningham, A.S.G., Salamini-Montemurri, M., Yang, J.H., Munk, R., et al. (2026). SenCat: Cataloging human cell senescence through multi-omic profiling of multiple senescent primary cell types. Mol Cell. 10.1016/j.molcel.2026.05.017.

58. Gurkar, A.U., Gerencser, A.A., Mora, A.L., Nelson, A.C., Zhang, A.R., Lagnado, A.B., Enninful, A., Benz, C., Furman, D., Beaulieu, D., et al. (2023). Spatial mapping of cellular senescence: emerging challenges and opportunities. Nat Aging 3, 776–790. 10.1038/s43587-023-00446-6.

