## Supplementary Figures for "Replicative senescence of neural progenitors induces astrocyte senescence in 2D cultures and human midbrain organoids"

#### Supplementary Figure 1

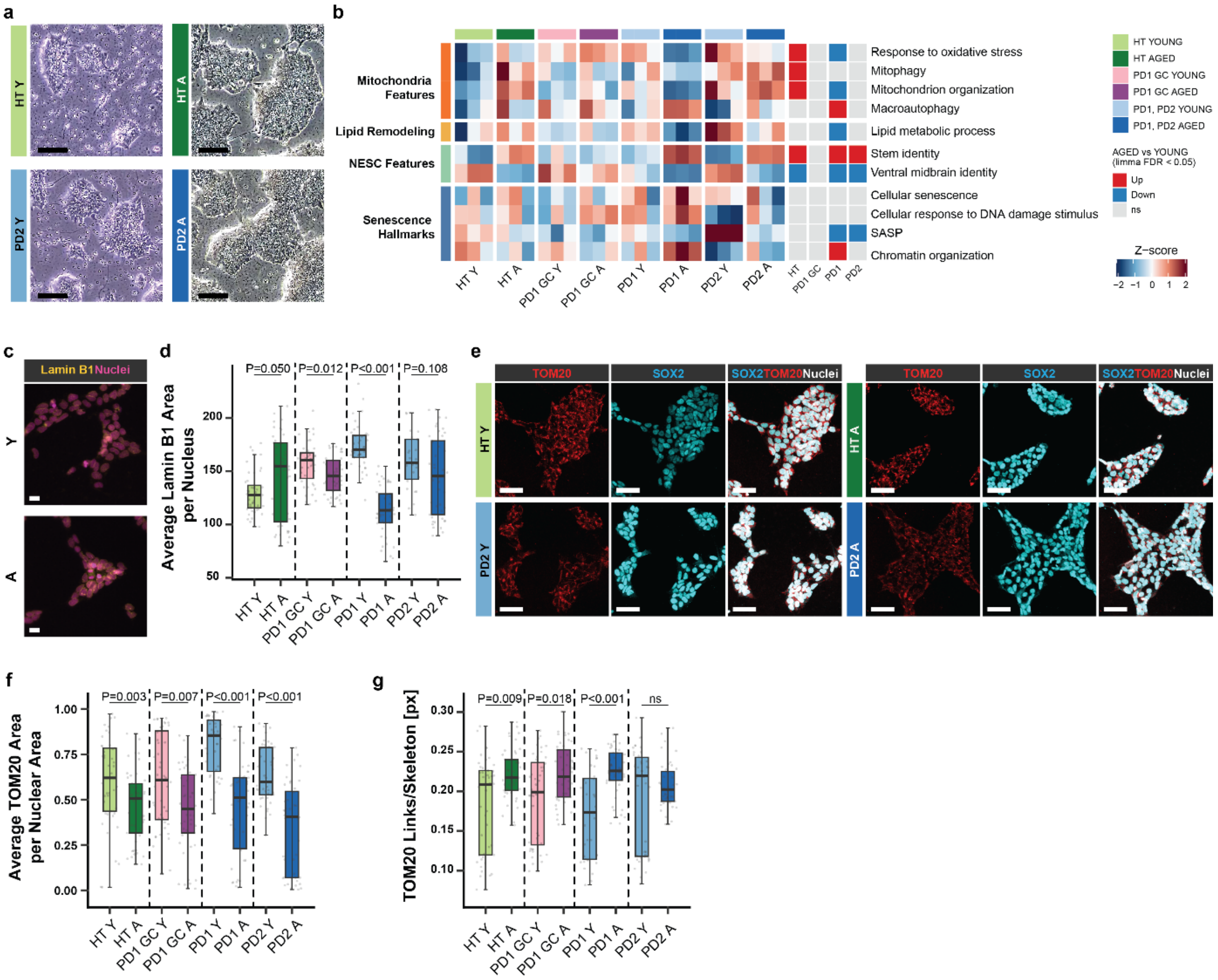


**Supplementary Figure 1: Extensive passaging induces nuclear lamina, and mitochondrial features alteration in NESCs.** (**a**) Representative brightfield images of NESC colonies. Scale bars, 100 µm. (**b**) Heatmap scoring of Mitochondria Features, Lipid Remodelling, NESC Features and Senescence Hallmarks selected, showing gene sets in Y and A NESCs. Scores were computed using the single-sample Gene Set Enrichment Analysis (ssGSEA) R package (Hanzelmann, Castelo et al. 2013). Blue, red, and grey squares indicate differential scores in pairwise Y versus A comparisons within each genetic background. (**d**) Quantification of mean Lamin B1 immunofluorescence signal area per nucleus. (**e**) Representative immunofluorescence images of the outer mitochondrial membrane marker TOM20. Single channel: TOM20 (red), SOX2 (cyan). Merged image includes TOM20 (red), SOX2 (cyan), and Nuclei (DAPI, grey). (**f, g**) Quantification of mean TOM20 positive area normalised per nuclear area (**f**) and mitochondrial branching quantified as the number of skeleton links normalised to mitochondrial skeleton (**g**). Scale bars, 50 µm.

For immunofluorescence-based analyses, three independent NESC batches per condition were analysed (Y: passages 5-7; A: passages 19-21). Each dot represents an individual image acquisition. HT, healthy control; PD1 and PD2, Parkinson's disease lines 1 and 2; PD1 GC, gene-corrected isogenic control of PD1 line; Y, young cells (<8 passages); A, aged cells (>15 passages). Mann-Whitney U tests were used for pairwise comparisons between Y and A within each genetic background in (d), (f), and (g).

#### Supplementary Figure2

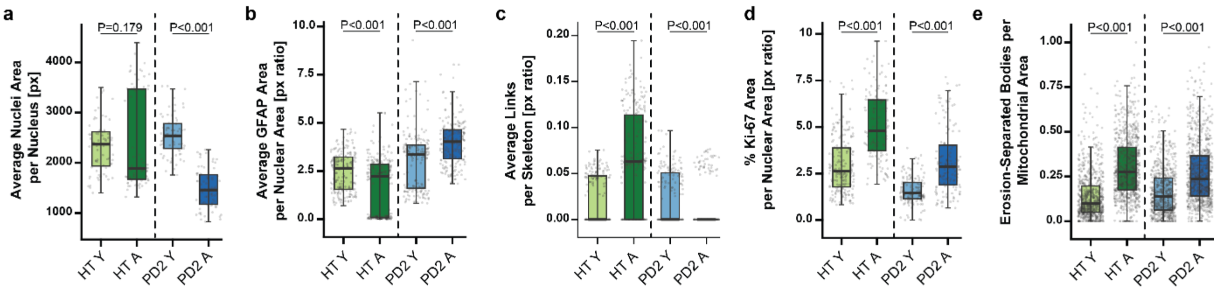

**Supplementary Figure 2: Extensive passaging of NSCs induces line-dependent senescence-like features in astrocytes while preserving cellular identity.** (**a**) Quantification of mean nuclear area per cell. (**b**) Quantification of GFAP area is normalised to nuclear area. (**c**) Quantification of astrocyte branching in immunofluorescence images, expressed as GFAP links normalized to GFAP skeleton area. (**d**) Percentage of Ki-67 positive area. Percentages for each data point are expressed as total Ki-67-positive area over the total nuclear area (**e**) Mitochondrial structural fragmentation quantified as erosion-separated mitochondrial bodies per mitochondrial area.

Three independent astrocyte batches per condition were analysed (Y: astrocytes derived from NSC passages 3-5; A: astrocytes derived from NSC passages 14-16). Each dot represents one independent image acquisition. HT, healthy control; PD2, Parkinson's disease line 2. Mann-Whitney U tests were used for pairwise comparisons between Y and A within each cell line, with P values indicated.

#### Supplementary Figure3

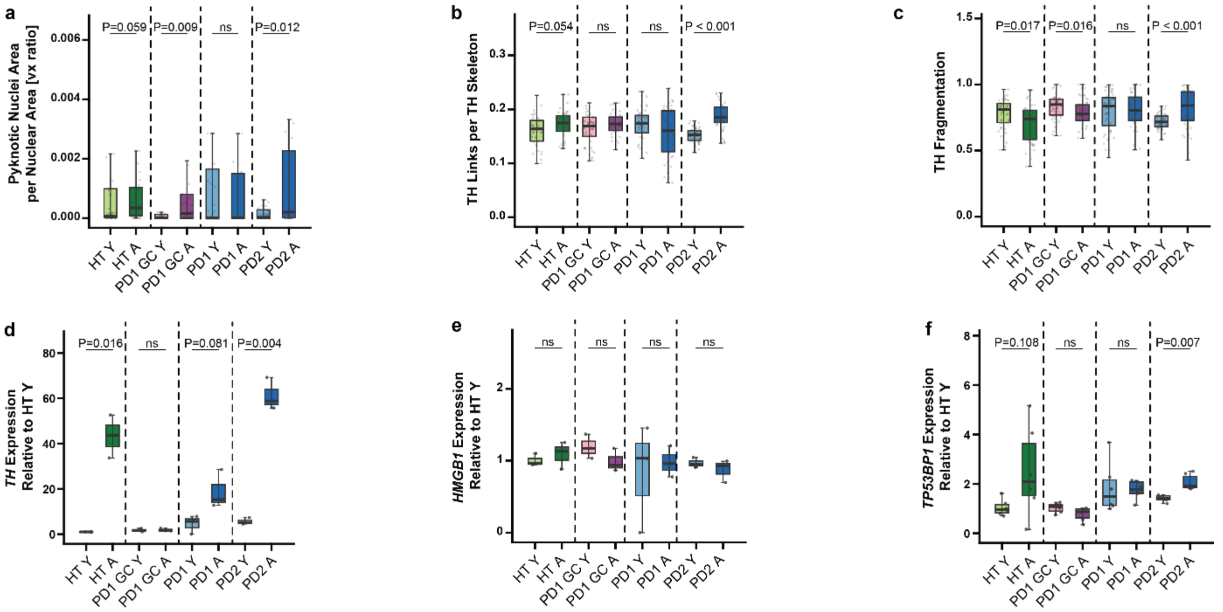


**Supplementary Figure 3: Extensive passaging of NESCs induces line-dependent senescence-like features in hMOs.** (**a**) Pyknotic nuclei are normalised to nuclear area. (**b**) Dopaminergic neuron branching was quantified as the number of skeleton links normalised to TH skeleton. (**c**) TH fragmentation quantification. (**d**) TH mRNA expression normalised to RPL37A and expressed relative to HT Y. (**e**) HMGB1 mRNA expression normalised to RPL37A and expressed relative to HT Y. (**f**) TP53BP1 mRNA expression normalised to RPL37A and expressed relative to HT Y.

For qPCR analysis, three independent hMO batches per condition were analysed (Y: passages 5-7; A: passages 17-19), each comprising pooled samples from at least 8 organoids. Two-sided t-tests were used for mRNA analyses. HT, healthy control; PD1 and PD2, Parkinson's disease lines; PD1 GC, gene-corrected isogenic control of PD1.

### Supplementary Figure4

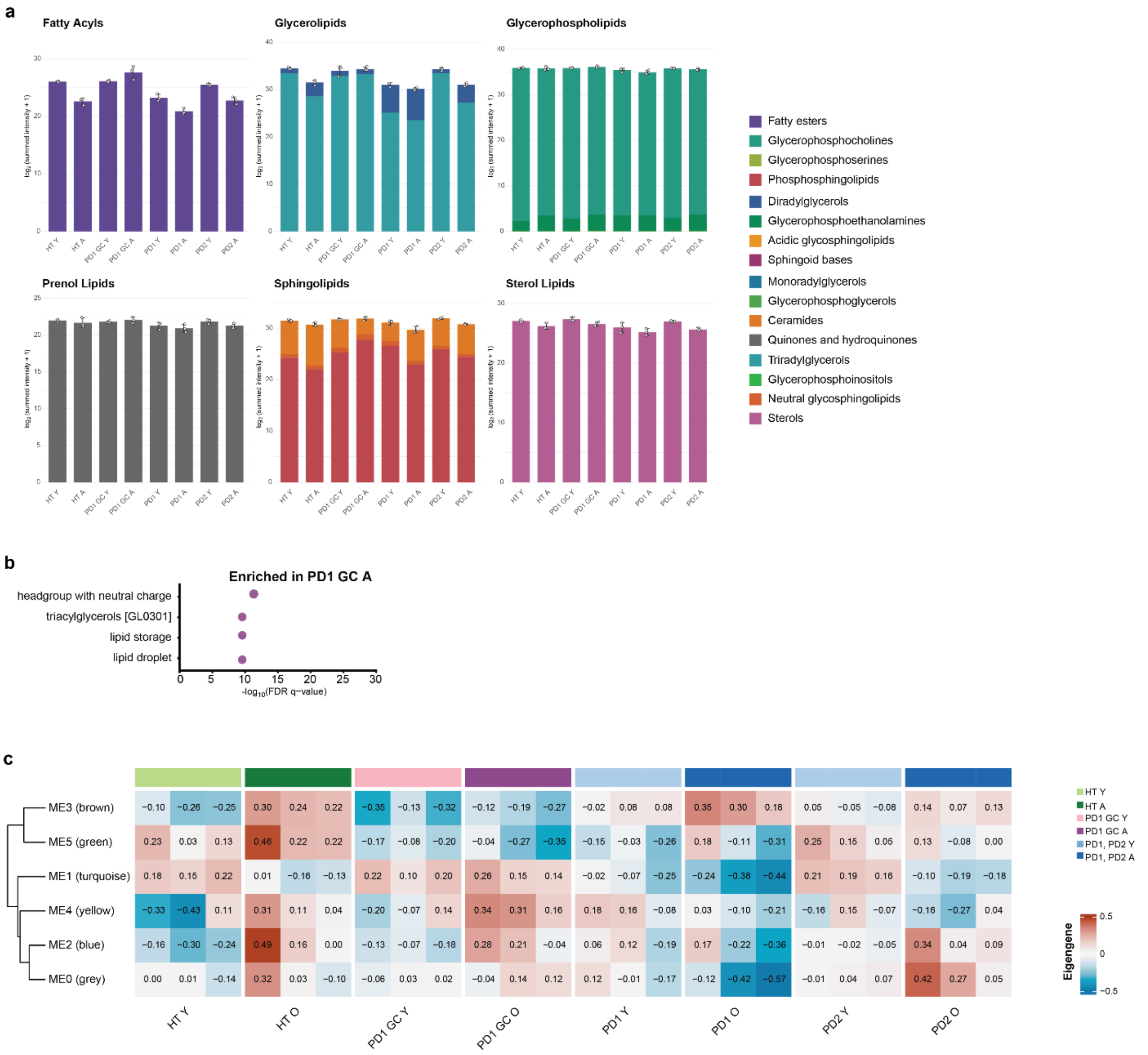

d

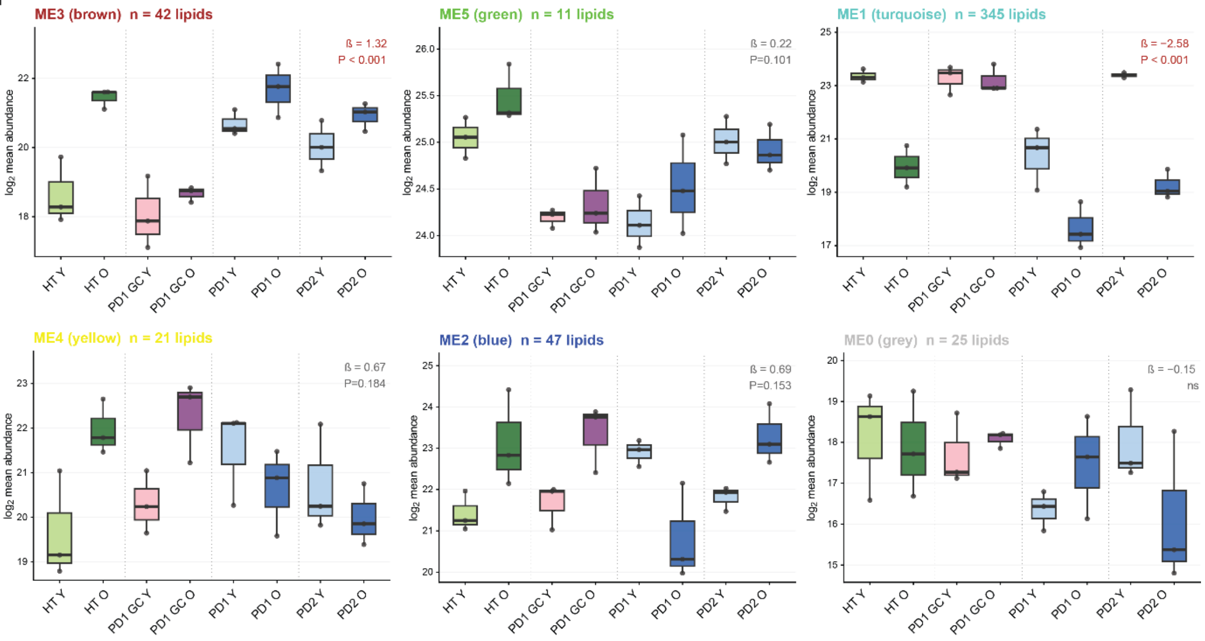


e

Module-Level Linear Model Results

Model: ME ~ Line * Aging

| Module | Aging β | Aging FDR | Interaction FDR | $R^2$ |
|---|---|---|---|---|
| ME3 | 0.450 | 1.16e-05 | 0.0156 | 0.908 |
| ME1 | −0.278 | 1.21e-03 | 0.0156 | 0.906 |
| ME2 | 0.450 | 9.13e-03 | 0.0624 | 0.553 |
| ME4 | 0.367 | 2.58e-02 | 0.0343 | 0.542 |
| ME5 | 0.171 | 1.78e-01 | 0.2530 | 0.696 |
| ME0 | 0.125 | 3.18e-01 | 0.0343 | 0.646 |

**Supplementary Figure 4: Extensive passaging of NESCs induces line-dependent lipidome changes in hMOs.** (**a**) Bar plot showing, for each sample, the summed lipid abundances per lipid category after log2(x+1) transformation. Colours indicate the contribution of individual lipid classes within each lipid category. (**b**) Lipid Ontology (LION) enrichment analysis of differentially abundant lipid species in young (Y) versus aged (A) PD1 GC hMOs. The x-axis indicates log10(FDR q-value); dot size represents the number of lipid species per enriched term. (**c**) Heatmap displaying module eigengene (ME) values for each sample derived from WGCNA analysis. Red indicates higher eigengene values and blue lower values relative to the module scale. (**d**) Boxplots showing the mean log2(x+1) lipid abundance of all lipid species assigned to each module, calculated per sample. Each dot represents one independent lipidomic sample. Statistical comparisons between Y and A conditions within each genetic background were performed using two-sided Wilcoxon rank-sum tests. Pearson correlation coefficients (**d**) between module mean abundance and the ageing variable are indicated. (**e**) Summary table of linear model results for module eigengenes (Model: ME ~ Line × Aging). The ageing coefficient (β) reflects the direction and magnitude of the ageing effect across genetic backgrounds, adjusted for line. Interaction FDR indicates genotype-dependent modulation of the ageing effect. P values were adjusted using the Benjamini-Hochberg method.

Three independent hMO batches per condition were analysed (Y: passages 5-7; A: passages 17-19), each comprising pooled samples from 5-8 organoids. Differential lipid abundance was assessed using linear modelling with Benjamini-Hochberg correction. Module-level statistics were derived from linear models (ME ~ Line × Aging) with Benjamini-Hochberg adjustment. HT, healthy control; PD1 and PD2, Parkinson's disease lines; PD1 GC, gene-corrected isogenic control.

#### Supplementary Figure5

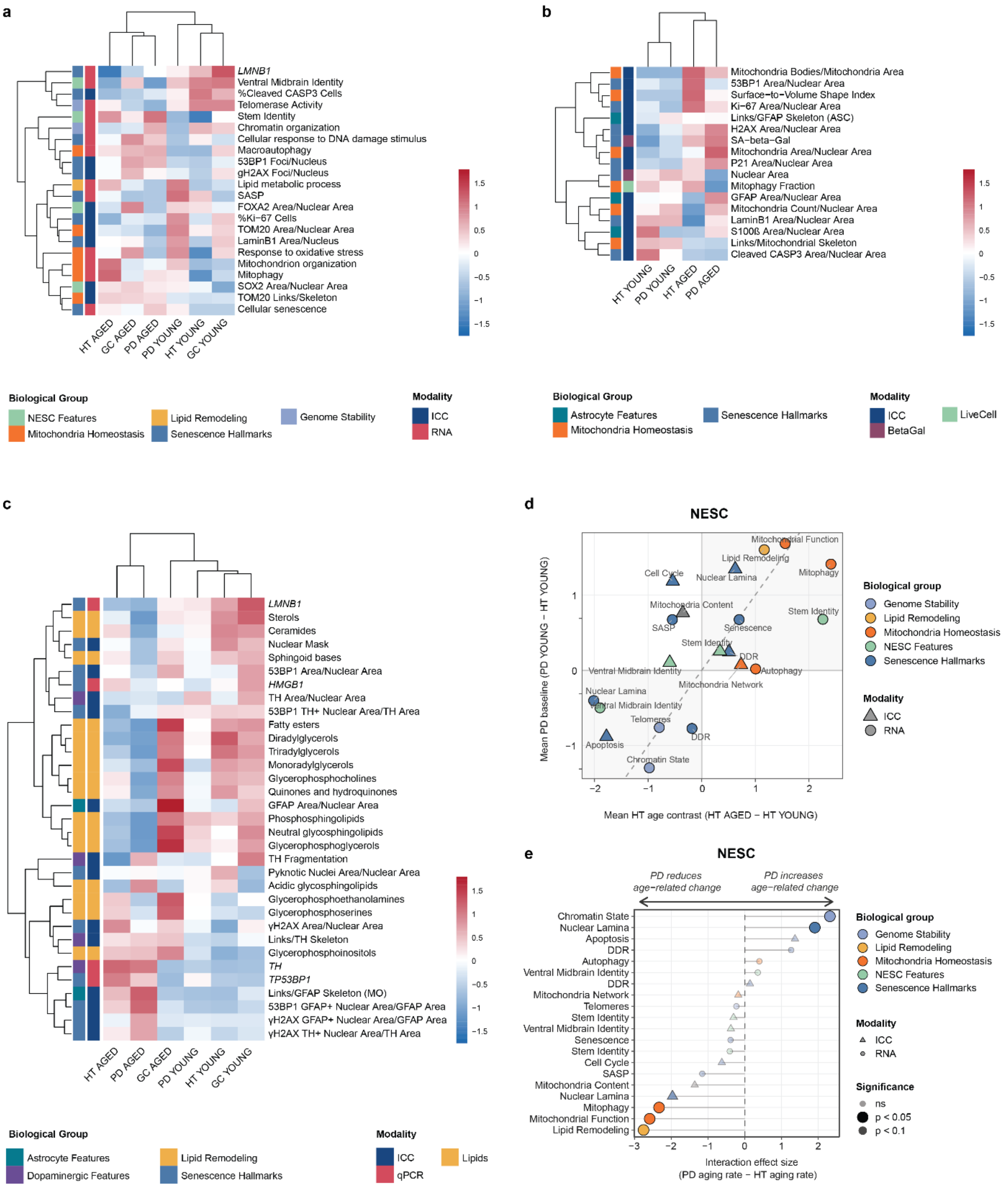


**Supplementary Figure 5: Category-level summary of senescence and PD-associated effects in NESCs.** (**a-c**) Heatmaps showing all individual biological features measured in NESC (**a**), astrocytes (**b**), and human midbrain hMOs (**c**). Each row represents a single feature, grouped by modality and macro-category (left annotation bars). Columns represent the six experimental conditions defined by genotype (HT, GC, PD) and age (YOUNG, AGED);

astrocyte analyses include only HT and PD lines. Each cell shows the mean z-score across biological replicates for that feature in that condition. PD1 and PD2 cell lines were analysed together. (**d**) Scatter plot of mean PD baseline effect (PD Y − HT Y; y-axis) versus mean HT ageing effect (HT A − HT Y; x-axis) at the biological category level in NESCs. Each point represents one biological category; colour indicates biological group, and point shape indicates measurement modality. The grey regions highlight features that show concordant behaviour in PD Y and HT A relative to the HT Y baseline. The dashed diagonal represents equal ageing and PD baseline effects. (**e**) Lollipop plot showing the interaction effect between genotype (PD vs. HT) and age on biological category scores in NESCs. The interaction effect quantifies the degree to which PD amplifies or attenuates age-related change (PD ageing rate - HT ageing rate). Each point represents one biological category × modality; colour indicates biological group and shape indicates measurement modality. Point transparency and size reflect statistical significance (opaque/large: $p < 0.05$; intermediate: $p < 0.1$; faded/small: not significant). Horizontal lines connect each category to the zero reference. Statistical significance was assessed using linear mixed-effects models with genotype × age interaction terms; p-values are uncorrected.

HT, healthy; PD, Parkinson's disease; GC, gene-corrected of PD1; Y, young (low passage); A, aged (high passage); ICC, immunocytochemistry-derived morphometric features; qPCR, mRNA expression; Lipids, lipidomic features.

### Supplementary Table 1. Materials and Equipment Table

| Primary Antibodies | | |
|---|---|---|
| **Antibody Name** | **Supplier** | **Catalog Number** |
| 53BP1 (rabbit polyclonal; 1:200) | NovusBio | Cat#NB100-304; RRID:AB_10003037 |
| Beta III Tubulin (chicken polyclonal; 1:1000) | Millipore (Merck) | Cat#AB9354; RRID:AB_570918 |
| Cleaved Caspase-3, Asp175 (rabbit; 1:200) | Cell Signaling Technology | Cat#9661; RRID:AB_2341188 |
| Glial Fibrillary Acidic Protein, GFAP (chicken polyclonal; 1:1000) | Millipore (Merck) | Cat#AB5541; RRID:AB_177521 |
| Histone H2AX (rabbit polyclonal; 1:200; astrocytes only | Sigma-Aldrich | Cat#SAB4501369; RRID:AB_10746039 |
| HNF-3beta, FoxA2, clone RY-7 (mouse monoclonal; 1:300) | Santa Cruz Biotechnology | Cat#sc-101060; RRID:AB_1124660 |
| Ki-67, clone B56 (mouse monoclonal; 1:200) | BD Biosciences | Cat#550609; RRID:AB_393778 |
| Lamin B1 (rabbit polyclonal; 1:500) | Abcam | Cat#ab16048; RRID:AB_443298 |
| Nestin, clone 10C2 (mouse monoclonal; 1:200) | Millipore (Merck) | Cat#MAB5326; RRID:AB_2251134 |
| p21 Waf1/Cip1, clone DCS60 (mouse monoclonal; 1:200) | Cell Signaling Technology | Cat#2946; RRID:AB_2260325 |
| Pax6 (rabbit polyclonal; 1:300) | Covance | Cat#901301; RRID:AB_291612 |
| Phospho-Histone H2AX (Ser139), γH2AX (rabbit polyclonal; 1:200; NESC only) | Cell Signaling Technology | Cat#2577; RRID:AB_2118010 |
| S100B (mouse monoclonal; 1:600) | Sigma-Aldrich | Cat#S2532; RRID:AB_477499 |
| SOX2 (rabbit polyclonal; 1:100) | Abcam | Cat#ab97959; RRID:AB_2341193 |
| TH, Tyrosine Hydroxylase (mouse monoclonal; 1:50) | Millipore (Merck) | Cat#MAB318; RRID:AB_2201528 |
| Tom20, clone F-10 (mouse monoclonal; 1:50) | Santa Cruz Biotechnology | Cat# sc-17764; RRID:AB_628381 |
| **Secondary Antibodies** | | |
| **Antibody Name** | **Supplier** | **Catalog Number** |
| DAPI (4',6-Diamidino-2-phenylindole dihydrochloride; 1 µg/mL) | Millipore (Merck) | Cat#D8417 |
| Donkey anti-Chicken IgY (H+L) Highly Cross Adsorbed, Alexa Fluor™ 647 (1:500) | Thermo Fisher / Invitrogen | Cat#A78952 |
| Donkey anti-Goat IgG (H+L) Cross-Adsorbed, Alexa Fluor 488/568/647 (1:1000) | Thermo Fisher / Invitrogen | Cat#A-11055/A-11057/A-21447 |
| Donkey anti-Mouse IgG (H+L) Highly Cross-Adsorbed, Alexa Fluor™ Plus 488/568/647 (1:500) | Thermo Fisher / Invitrogen | Cat# A-21202/A32787/ A-31571 |
| Donkey anti-Rabbit IgG (H+L) Highly Cross-Adsorbed, Alexa Fluor™ Plus 488/568/647 (1:500) | Thermo Fisher / Invitrogen | Cat# A-21206/ A10042 /A-31573 |
| Goat anti-Chicken IgY (H+L), Alexa Fluor 488/568/647 (1:1000) | Thermo Fisher / Invitrogen | Cat#A-11039/A-11041/A-21449 |
| Goat anti-Mouse IgG (H+L) Cross-Adsorbed, Alexa Fluor 488/568/647 (1:1000) | Thermo Fisher / Invitrogen | Cat#A-11001/A-11004/A-21235 |
| Goat anti-Rabbit IgG (H+L) Cross-Adsorbed, Alexa Fluor 488/568/647 (1:1000) | Thermo Fisher / Invitrogen | Cat#A-11008/A-21244/A-21245 |
| Hoechst 33342 (20 mM; 1:1000; astrocytes only) | Thermo Fisher / Invitrogen | Cat#62249 |
| **Chemicals, peptides, and recombinant proteins** | | |
| **Product Name** | **Supplier** | **Catalog Number** |
| Accutase cell dissociation reagent | Thermo Fisher | Cat#A11105-01 |
| B-27 Supplement (50×, with Vitamin A) | Gibco (Thermo Fisher) | Cat#17504044 |
| B-27 Supplement without Vitamin A (50×) | Gibco (Thermo Fisher) | Cat#12587010 |
| BSA (Bovine Serum Albumin) | Sigma-Aldrich | Cat#A4503 |
| CHIR99021 (GSK3β inhibitor) | Axon Medchem | Cat#CT99021 |
| Chloroform | Sigma-Aldrich | Cat#319988 |
| Dibutyryl-cAMP | BioSynth | Cat#ND07996 |
| DMEM/F-12 (1:1) medium | Gibco (Thermo Fisher) | Cat#11320033 |
| DMSO (dimethyl sulfoxide) | Sigma-Aldrich | Cat#D8418 |
| Dorsomorphin (BMP inhibitor) | Tocris Bioscience | Cat#3093 |
| Essential 8 (E8) medium | Gibco (Thermo Fisher) | Cat#A1517001 |
| Essential 8 Flex medium | Gibco (Thermo Fisher) | Cat#A2858501 |
| Fetal Bovine Serum (FBS) | Sigma-Aldrich | Cat#F7524 |
| Fluoromount-G aqueous mounting medium | Thermo Fisher | Cat#00-4958-02 |
| Geltrex reduced growth factor basement membrane matrix | Gibco (Thermo Fisher) | Cat#A1413302 |

| | | |
|---|---|---|
| GlutaMax supplement (100×) | Gibco (Thermo Fisher) | Cat#35050061 |
| KnockOut Serum Replacement | Gibco (Thermo Fisher) | Cat#10828010 |
| KO-DMEM (Knockout DMEM) | Gibco (Thermo Fisher) | Cat#10829018 |
| L-Ascorbic acid | Sigma-Aldrich | Cat#A4544 |
| L-Glutamine | Tocris Bioscience | Cat#0218 |
| Low-melting-point agarose (3%) | Biozym Scientific | Cat#840100 |
| MEM Non-Essential Amino Acids (NEAA, 100×) | Gibco (Thermo Fisher) | Cat#11140050 |
| N-2 Supplement (100×) | Gibco (Thermo Fisher) | Cat#17502001 |
| Neurobasal medium | Gibco (Thermo Fisher) | Cat#21103049 |
| Normal Donkey Serum (NDS) | Abcam | Cat#ab7475 |
| Normal Goat Serum (NGS) | Abcam | Cat# ab138478 |
| Paraformaldehyde (PFA) 4% | Millipore (Merck) | Cat#1.00496.5000 |
| PBS-Tween (Tween-20 solution) | Applichem | Cat#A1389,0500 |
| Penicillin/Streptomycin (100×) | Gibco (Thermo Fisher) | Cat#15140122 |
| Phosphate Buffered Saline (PBS, 1×) | Thermo Fisher | Cat#10010023 |
| Purmorphamine (PMA, Smoothened agonist) | Enzo Life Sciences | Cat#ALX-420-045-M005 |
| Recombinant human BDNF | PeproTech | Cat#450-02 |
| Recombinant human EGF | PeproTech | Cat#100-20 |
| Recombinant human FGF2 (basic FGF) | PeproTech | Cat#100-15 |
| Recombinant human GDNF | PeproTech | Cat#450-10 |
| Recombinant human LIF | Sigma-Aldrich | Cat#L5283 |
| Recombinant human TGF-β3 | Thermo Fisher | Cat#100-36E |
| SB-431542 (TGF-β / ALK5 inhibitor) | Abcam | Cat#ab120163 |
| Sodium azide | Carl Roth | Cat#K305.1 |
| Triton X-100 | Carl Roth | Cat#3051.3 |
| TRIzol reagent | Thermo Fisher | Cat#PR94757 |
| U-bottom ultra-low attachment 96-well plate | faCellitate | Cat#F202003 |
| Y-27632 ROCK inhibitor | Millipore (Merck) | Cat#SCM075 |
| β-mercaptoethanol | Gibco (Thermo Fisher) | Cat#21985023 |

**Commercial kits**

| Product Name | Supplier | Catalog Number |
|---|---|---|
| AccuraCode RNA-Seq Kit V2 | Singleron Biotechnologies GmbH | Cat#10710174 |
| DNase I, amplification grade | Sigma-Aldrich (Merck) | Cat#AMPD1 |
| High Sensitivity RNA ScreenTape System | Agilent Technologies | Cat#5067-5579 |
| High-Capacity RNA-to-cDNA™ Kit | Applied Biosystems (Thermo Fisher) | Cat#4387406 |
| iQ SYBR Green Supermix | Bio-Rad | Cat#1708880 |
| iScript cDNA Synthesis Kit | Bio-Rad | Cat#1708890 |
| Maxwell RSC simplyRNA Tissue Kit | Promega | Cat#AS1340 |
| Phasemaker tubes | Thermo Fisher | Cat#15645268 |
| Qubit RNA High Sensitivity (HS) Assay Kit | Invitrogen (Thermo Fisher) | Cat#Q10211 |
| RNeasy Mini Kit (RNA purification) | Qiagen | Cat#74106 |
| Senescence Detection Kit (β-galactosidase activity) | Abcam | Cat#ab65351 |
| TaqMan Universal PCR Master Mix II, no UNG | Applied Biosystems (Thermo Fisher) | Cat#4440040 |

**Primers**

| Gene Name | Source | Forward | Reverse |
|---|---|---|---|
| *HMGB1* | Eurogentec | GCGAAGAAACTGGGAGAGATGTG | GCATCAGGCTTTCCTTTAGCTCG |
| *LMNB1* | Eurogentec | GTATGAAGAGGAGATTAACGAGAC | TACTCAATTTGACGCCCAG |
| *RPL37A* | Sigma-Aldrich | GTGGTTCCTGCATGAAGACAGTG | TTCTGATGGCGGACTTTACCG |
| *TH* | | AGTACACCGCCGAGGAGATT | GTGGCGGATATACTGGGTGC |

**TaqMan Primers**

| Gene Name | Source | Assay ID | Catalog Number |
|---|---|---|---|
| *P16/CDKN2A* | Thermo Fisher | Hs00923894_m1 | Cat#4331182 |
| *RPL37A* | Thermo Fisher | HS03044965_g1 | Cat#4331182 |
| *TP53BP1* | Thermo Fisher | Hs00996827_m1 | Cat#4331182 |

**Other Equipment**

| Product Name | Supplier | Catalog Number |
|---|---|---|
| AggreWell 800 24-well plate (embryoid body formation) | StemCell Technologies | Cat#27845 |
| U-bottom ultra-low attachment 96-well plate (hMO generation) | faCellitate | Cat#F202003 |

**Human iPSCs**

| Cell Line Name | Acronym | Internal Identifier | Gene correction/ Gene editing | Genetic Background for *LRRK2* | Age of Sampling | Age of onset | Sex | References |
|---|---|---|---|---|---|---|---|---|
| HT | A13777 | GMO168 | no | WT | Newborn | - | F | (Burridge, Thompson et al. 2011) |
| HT | A13777 Mitophagy | GMO170 | yes, mitophagy reporter insertion (Arias-Fuenzalida, Jarazo et al. 2019) | WT | Newborn | - | F | (Burridge, Thompson et al. 2011) |
| PD1 | T413MUT | GMO161 | no | LRRK2-G2019S | 51 | 40 | F | (Reinhardt, Glatza et al. 2013, Reinhardt, Schmid et al. 2013) |
| PD1 GC | T413GC | GMO162 | yes, correction of the LRRK2 point mutation | WT (gene corrected) | 51 | 40 | F | (Reinhardt, Glatza et al. 2013, Reinhardt, Schmid et al. 2013) |
| PD2 | 33879 | GMO169 | no | LRRK2-G2019S | 66 | 46 | F | (Nickels, Walter et al. 2019, Walter, Bolognin et al. 2019) |
| PD2 | 33879 Mitophagy | GMO171 | yes, mitophagy reporter insertion (Arias-Fuenzalida, Jarazo et al. 2019) | LRRK2-G2019S | 66 | 46 | F | (Arias-Fuenzalida, Jarazo et al. 2019) |
